# Representational dynamics during extinction of fear memories in the human brain

**DOI:** 10.1101/2025.04.26.650560

**Authors:** Daniel Pacheco-Estefan, Antoine Bouyeure, George Jacobs, Marie-Christin Fellner, Katia Lehongre, Virginie Lambrecq, Valerio Frazzini, Vincent Navarro, Onur Güntürkün, Lu Shen, Jing Yang, Biao Han, Qi Chen, Nikolai Axmacher

## Abstract

Extinction learning – the suppression of a previously acquired fear response – is critical for adaptive behavior and core for understanding the etiology and treatment of anxiety disorders. Electrophysiological studies in rodents have revealed critical roles of theta (4-12Hz) oscillations in amygdala and hippocampus during both fear learning and extinction, and engram research has shown that extinction relies on the formation of novel, highly context-dependent memory traces that suppress the initial fear memories. Whether similar processes occur in humans and how they relate to previously described neural mechanisms of episodic memory formation and retrieval remains unknown. Intracranial EEG (iEEG) recordings in epilepsy patients provide direct access to the deep brain structures of the fear and extinction network, while representational similarity analysis (RSA) allows characterizing the memory traces of specific cues and contexts. Here we combined these methods to show that amygdala theta oscillations during extinction learning signal safety rather than threat and that extinction memory traces are characterized by stable and context-specific neural representations that are coordinated across the extinction network. We further demonstrate that context specificity during extinction learning predicts the reoccurrence of fear memory traces during a subsequent test period, while reoccurrence of extinction memory traces predicts safety responses. Our results reveal the neurophysiological mechanisms and representational characteristics of context-dependent extinction learning in the human brain. In addition, they show that the mutual competition of fear and extinction memory traces provides a mechanistic basis for clinically important phenomena such as fear renewal and extinction retrieval.

## Introduction

In a constantly changing world, the ability to update previously learned knowledge is essential for adaptive behavior. The neural mechanisms that support this ability have been extensively studied within the domain of extinction learning. Extinction learning allows for the inhibition of previously acquired unconditioned responses (in particular, fear responses) that have become dysfunctional; accordingly, deficits of extinction learning are associated with numerous psychiatric conditions, most prominently anxiety disorders and depression, and are targeted during exposure therapy (Beckers et al., 2023; Bouton et al., 2021; Maren et al., 2013; McNally, 2007).

While early accounts of extinction learning suggested that it weakens the association between a conditioned stimulus (CS) and an unconditioned stimulus (US) (Rescorla & Wagner, 1972), more recent theories propose that it consists in the formation of a new inhibitory memory trace that suppresses the initial fear response (Bouton, 2004; Y. Liu et al., 2024; Maren et al., 2013; Maren & Quirk, 2004). This new memory trace is highly context-dependent, as demonstrated by the phenomenon of renewal, where fear responses re-emerge when tested either in the initial (acquisition) context or in a new context (Bouton, 2004; Corcoran & Maren, 2004; Maren et al., 2013; Orsini et al., 2011; Orsini & Maren, 2012; Quirk & Mueller, 2008).

Human neuroimaging studies have provided a detailed understanding of the circuits involved in fear learning and extinction, identifying a core network that includes amygdala (AMY), hippocampus (HPC), and regions of prefrontal cortex (PFC; Fullana et al., 2016, 2018). Among these regions, PFC and HPC appear particularly relevant for the context specificity of extinction learning (Bouton, 2004; Eichenbaum, 2017a, 2017b; Gilmartin et al., 2014; Maren et al., 2013; Maren & Quirk, 2004; Orsini et al., 2011). Indeed, the PFC seems to be more involved in the inhibition of fear responses (Gilmartin et al., 2014; Maren et al., 2013) and in the contextual regulation of fear during and after extinction (Orsini et al., 2011) than in the acquisition of novel fear memories. Furthermore, recent studies indicate that the lateral PFC (lPFC) supports flexible, task-dependent updating of memory traces (Pacheco-Estefan et al., 2024), consistent with its role in adaptive cognitive control (Darna et al., 2024; Weber et al., 2023).

Research in rodents has provided important insights into the neural mechanisms of fear acquisition, expression and extinction. Notably, several studies have shown that the amplitude of theta (4-12Hz) oscillations in the prelimbic cortex, dorsal anterior cingulate cortex (dACC), and/or amygdala increases during fear acquisition (Paré & Collins, 2000) and expression (Courtin et al., 2014; Fenton et al., 2014; Karalis et al., 2016; Lesting et al., 2011; Likhtik et al., 2014; Rahman et al., 2018; Seidenbecher et al., 2003; Taub et al., 2018). Similarly, human EEG studies have reported heightened frontocentral theta oscillations, particularly during fear conditioning (e.g., Chen et al., 2021; Pirazzini et al., 2023) and fear recall (e.g., Bierwirth et al., 2021; Mueller et al., 2014; Sperl et al., 2019; but see (Bierwirth et al., 2023). In addition, studies in rodents have shown that long range theta (2-12Hz) connectivity across regions of the extinction network, including AMY, PFC and HPC, has been associated with the discrimination of aversive and safe cues (Likhtik et al., 2014), and with the extinction of fear associations (Lesting et al., 2011, 2013; Nguyen et al., 2023).

In humans, the electrophysiological mechanisms underlying extinction learning in AMY and HPC are largely unknown because these regions are difficult to investigate non-invasively. Intracranial EEG (iEEG) recordings in epilepsy patients provide invaluable data about neurophysiological processes in deep brain areas at the highest temporal and spatial resolution possible (Axmacher, 2023; Parvizi & Kastner, 2018). Indeed, iEEG research in the domain of episodic memory has provided important insights into the mesoscopic neural mechanisms underlying memory formation and retrieval (Burke et al., 2014; Fell et al., 2011; Foster et al., 2012, 2015; Gattas et al., 2023; Lega et al., 2012; Miller et al., 2013; Seger et al., 2023), including evidence for a prominent role of theta oscillations (Herweg et al., 2020). Other recent iEEG studies described the neurophysiological mechanisms in AMY and HPC underlying the impact of aversive emotions on episodic memory formation and retrieval (Costa et al., 2022, 2024; Qasim et al., 2023; H. Zhang et al., 2024; Zheng et al., 2017).

Despite this progress in understanding episodic memory and its modulation by aversive emotions, surprisingly few iEEG studies explored the neurophysiological mechanisms of fear learning and extinction, and thus it is unclear whether and to which extent they rely on the same neural machinery that supports episodic memory. A notable exception is the study by Chen et al. (2021) who reported increased theta oscillations in AMY and dorsomedial PFC following aversive stimuli during fear learning. Whether AMY and PFC theta frequency oscillations play a role during extinction learning remains unknown.

In addition to neural oscillations, rodent research has described how memory traces of specific experiences – engrams – are built in AMY, HPC, and other areas during fear learning and how they are suppressed or modified during extinction (X. Liu et al., 2012; X. Zhang et al., 2020). In humans, Representational Similarity Analysis (RSA; Kriegeskorte et al., 2008; Kriegeskorte & Diedrichsen, 2019) is increasingly used to identify stimulus-specific memory traces, i.e. the neural representations of unique episodes and events, at a meso- and macroscopic level (Griffiths et al., 2019; J. Liu et al., 2021; Michelmann et al., 2016; Pacheco Estefan et al., 2019; Staresina et al., 2016; Yaffe et al., 2014). Applied to iEEG data, this approach typically relies on patterns of the power of neural oscillations across frequencies and electrodes (Manning, 2023; Pacheco Estefan, 2023; Xie et al., 2023). RSA can then either be used to calculate the similarity of representations between encoding and retrieval (reinstatement, or encoding-retrieval similarity) or to measure similarities between representations of different items during either encoding or retrieval.

Episodic memory studies using RSA revealed fundamental differences in the “representational formats” of memory traces of item-context associations as compared to single items: While memories of individual items depend on broad frequency ranges and on sensory regions – for visual stimuli, in lateral temporal cortex (TMP) – memory traces of item-context associations exhibit a strong reliance on theta frequency oscillations and the HPC (Pacheco Estefan et al., 2019; Staresina et al., 2016). Because of the strong context dependency of extinction learning, one may expect similar representational signatures as for episodic memory traces of item-context associations. Furthermore, RSA studies using functional magnetic resonance imaging (fMRI) demonstrated that the “stability” of item representations – i.e., the similarity of neural activity patterns across repeated exposures – correlates with successful memory formation (Lu et al., 2015; Xue et al., 2010). Similar effects were observed during fear acquisition, where higher levels of item stability predicted learning success (Visser et al., 2013). Here we investigated the neurophysiological mechanisms and representational characteristics of fear and extinction memory traces and their impact on subsequent renewal. We developed a novel fear and extinction learning paradigm that we conducted with epilepsy patients (N = 49) who were implanted with iEEG electrodes across the fear and extinction network (Figure 1A). During acquisition, patients were exposed to images of three electric devices, two of which were paired with an aversive stimulus (CS+) while the third one was not (CS-). During extinction, the contingency of one CS+ cue changed, resulting in three different types of cues: CS+_acquisition_/CS+_extinction_ (CS++), CS+_acquisition_/CS- _extinction_ (CS+-) and CS-_acquisition_/CS- _extinction_ (CS--) (Figure 1A, top). During the final test phase, none of the items was paired with a US. Critically, to study the specificity of context representations, the CS items were presented within four different thematically related context videos in each experimental phase (e.g., four snow landscape videos during acquisition, four videos during extinction, and four videos during test), corresponding to an “ABC” paradigm (Figure 1A, top). This paradigm allowed us to track the memory traces of items and contexts during fear acquisition, extinction, and renewal across key regions of the extinction network including AMY, HPC, and PFC regions, as well as sensory processing areas in TMP (Figure 1B).

**Figure 1.**
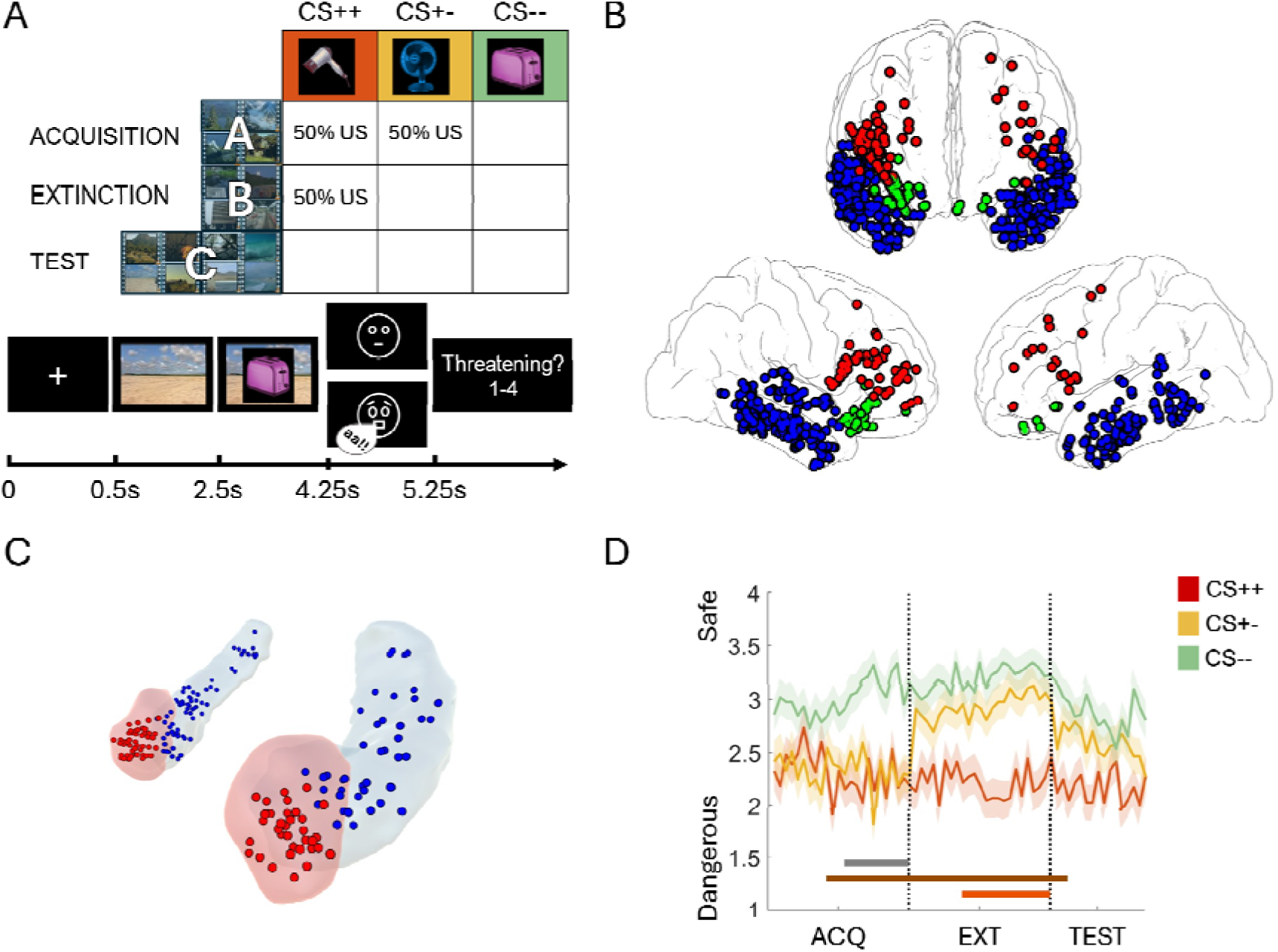
Experimental paradigm, electrode implantation and behavioral results. A) Experimental design. Top: Contingencies of consistently threatening CS++ cues (red), consistently safe CS--cues (green) and cues with changing contingencies (yellow) across the tree experimental phases (acquisition, extinction, and test). Cues were presented together with a set of context videos that were unrelated to the contingency in a given trial and that changed between phases (ABC paradigm). Bottom: Trial structure. Contexts were presented first, but remained on screen during CS presentation. The US consisted of either a neutral face (CS-trials) or a fearful face paired with a loud scream (CS+ trials). The cartoon faces shown are for illustrative purposes only and do not depict the actual stimuli used in the experiment. Participants rated the perceived cue contingencies on each trial. B) Electrode implantation. Selected contacts across our group of participants are overlaid on an average brain surface in Montreal Neurological Institute (MNI) coordinates. Neocortical regions of interest selected for analysis included electrodes implanted bilaterally in temporal (shown in blue), prefrontal (shown in red) and orbitofrontal (shown in green) cortices. C) Electrode locations in the amygdala (shown in red) and the hippocampus (shown in blue). D) Behavioral performance. Wilcoxon signed rank tests were conducted at every trial position to investigate the progressive learning of task contingencies across experimental phases. Regions where significant effects were observed after correction for multiple comparisons are indicated at the bottom of the figure (cluster-based permutation statistics with 1,000 permutations; grey: CS+- vs. CS--; brown: CS++ vs. CS--; orange: CS+- vs. CS++). Shaded lines depict group average ratings at each trial position ± S.E.M.

## Results

### Behavioral results

We first analyzed the behavioral responses to the different cue types across experimental phases. A two-way ANOVA with “cue type” (CS++, CS+-, CS--) and “experimental phase” (acquisition, extinction, test) as repeated measures revealed significant main effects of both cue type (F(2, 94) = 19.02; p = 1.16e-07) and experimental phase (F(2, 94) = 9.03; p = 2.58e-04) as well as a significant interaction (F(4, 188) = 6.96; p = 3.02e-05). Importantly, cues with changing contingency (CS+-) were perceived as significantly less threatening during extinction than during acquisition (t(47) = 5.67; p = 7.44e-06, Bonferroni corrected), confirming that participants correctly adjusted their expectations of the US.

We evaluated the progressive learning of trial contingencies during acquisition and extinction. We sorted trials according to their temporal position (i.e., trial number) separately in the three conditions (CS++, CS+-, and CS--), and compared responses at every position using paired Wilcoxon signed rank tests, followed by cluster-based permutation statistics to correct for multiple comparisons (Methods). During acquisition, ratings of CS++/CS+- vs. CS-- cues started to differ at trials 10 and 13, respectively (both p = 0.001; brown and grey lines in Figure 1C), while ratings did not differ between CS++ and CS+- items, which were both threatening during acquisition (red and yellow line, all p > 0.72). During extinction, ratings started to differ between CS++ and CS+- cues at trial 9 (p = 0.001), indicating that participants gradually learned the new contingencies, while they differed immediately between CS++ vs. CS-- cues (p = 0.001; Figure 1C, orange and brown line). Ratings of CS+- and CS-- cues did not differ significantly (all p > 0.46). During the test phase, ratings of CS++ and CS-- differed in the first 3 trials, while the remaining contrasts did not show any significant differences. However, comparisons of average ratings during the test revealed significant differences between CS++ and CS+- trials (t(47) = 2.51, p = 0.015) and between CS++ and CS-- trials (t(47) = 3.1, p = 0.003), whereas no significant difference was observed between CS+- and CS-- trials (t(47) = 1.13, p = 0.26). Taken together, these results show that participants accurately learned the task contingencies and their changes throughout the experiment. In addition, while differences in ratings diminished during the test phase, participants still rated CS++ items on average as more threatening than CS-- or CS+- items, indicating that fear responses did not completely disappear.

### Amygdala theta oscillations signal safety during extinction

Previous literature has established a key role of theta (4-12Hz) oscillations in AMY, HPC and PFC for fear learning and extinction, in both rodents (Seidenbecher et al., 2003) and humans (Chen et al., 2021). We thus focused on this frequency band in our initial analysis and compared differences in theta power between CS+ and CS- cues. Following previous studies (e.g., Chen et al., 2021), we specifically assessed the time period from cue onset until US presentation (in reinforced CS+ trials, or corresponding time point in the other trials), during both acquisition and extinction.

During acquisition, we found no significant power differences between CS+ and CS- cues in any of our regions of interests (ROIs; all p > 0.54; see Figure 2A, left for the results in the AMY). During extinction, theta power was significantly higher for CS- as compared to CS+ trials in the AMY (p_corr_ < 0.005; Figure 2A, right), while no significant power differences were observed in any other ROI (all p_corr_ > 0.24). This AMY effect occurred at a relatively late time period preceding the US presentation (i.e., from 1.18 to 1.75s) and was confined to the theta (3-10Hz) frequency range (Figure 2A, right). Follow up analyses revealed that theta power was significantly lower in CS++ trials compared to both CS+- (p = 0.005; Figure 2B, top) and CS-- trials (p = 0.026; Figure 2B, middle), with effects occurring in overlapping time frequency periods to those observed in the “current valence” contrast in Figure 2A, right. No significant differences were observed between CS+- and CS-- trials (p = 0.38; Figure 2B, bottom). To determine whether the difference between CS+ and CS- trials was specific to the theta band, we analyzed all frequencies across the 1-100Hz range. Results confirmed that differences between conditions were localized to the same cluster observed in Figure 2A, right, while no significant effects were observed in other time-frequency bins (p = 0.004; Figure 2C, top). Further analyses in this cluster revealed that effects were driven by theta power increases above baseline for both CS+- (t(31) = 3.21, p = 0.003) and CS-- items (t(31) = 2.3, p = 0.027), as well as theta power reductions below baseline for CS++ items (t(31) = -3.9, p = 0.0004; Figure 2C, bottom). As a baseline, we chose the average across all trials in all phases (see Methods). No differences between CS+ and CS- trials occurred in this time-frequency cluster during acquisition (t(31) = -0.69, p = 0.5). Please note that these latter analyses are not circular because the time-frequency cluster is defined by contrasting CS+ and CS- trials during extinction, and not by comparing the three different trial types (CS++, CS+-, CS--) to zero. Together, these results show increases of AMY theta power during extinction for CS- as compared to CS+ trials. They also reveal theta power in CS- trials during extinction were systematically above baseline, which may reflect a safety signal.

**Figure 2.**
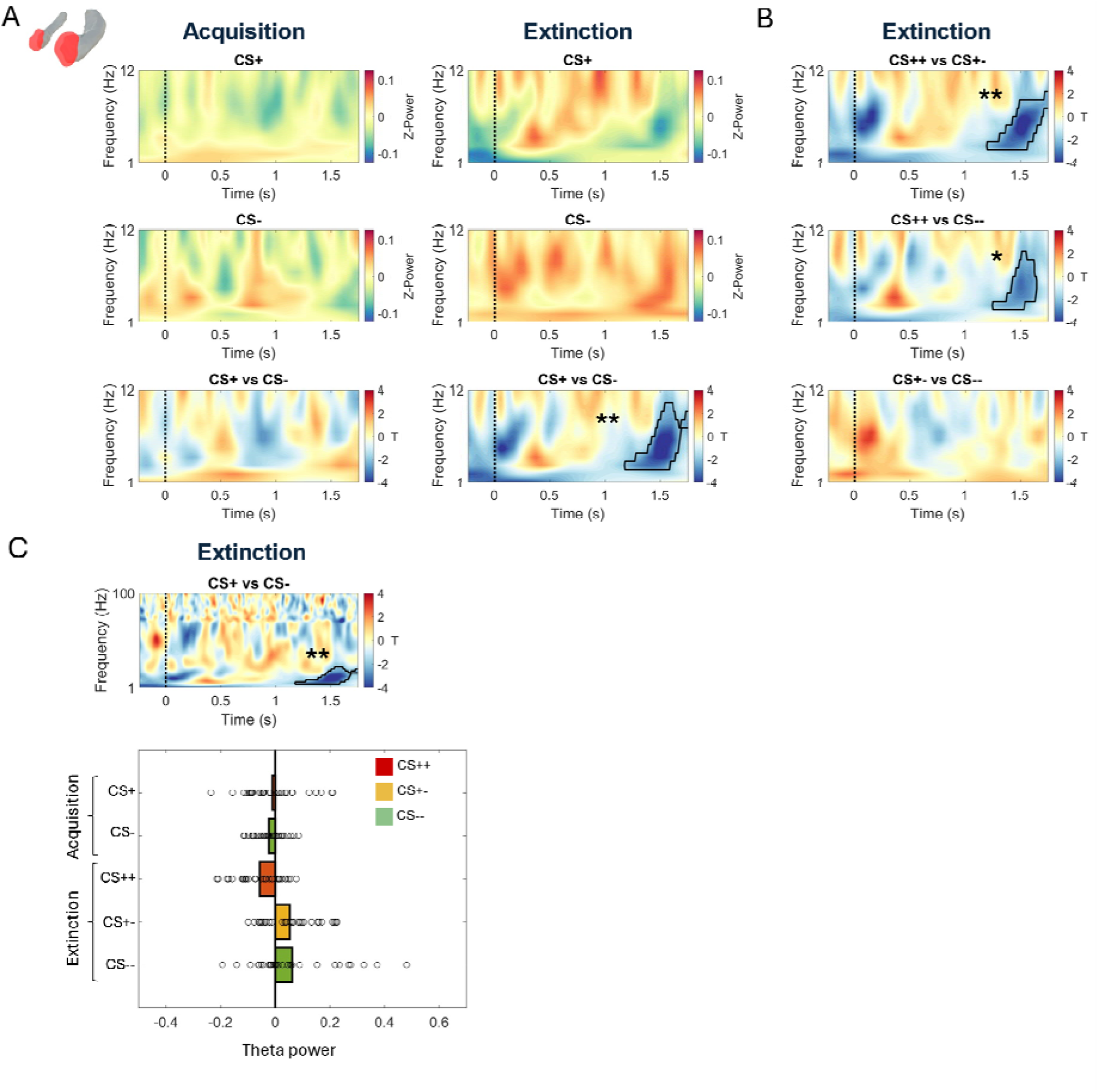
Higher theta power in the amygdala for CS- as compared to CS+ trials during extinction. A) Amygdala 1-12Hz power. During acquisition (left), no significant differences in low frequency power were observed between CS+ and CS- trials. Bottom row shows the result of the contrast between the two conditions, while top and middle rows reflect levels of Z-scored power for each condition. During extinction (right), oscillatory power was significantly higher for CS- trials in a late time period of cue acquisition, i.e., from 1.18 to 1.75s in the 3-10Hz frequency range. Negative T-values depicted in blue reflect higher theta power for CS- items. B) T-maps of the CS++ vs C+- (top), CS++ vs CS-- (middle) and CS+- vs CS-- (bottom) contrasts. C) Top: CS+ vs CS- contrast in the frequency spectrum between 1-100Hz. Bottom: average theta power in the cluster observed in panel A, bottom right, and C, top, for all CS types during the phases of acquisition and extinction. In panels A, B and C, significant regions surviving correction for multiple comparisons using cluster-based permutation statistics are outlined in black in the time-frequency maps. *: p < 0.05; **: p < 0.01.

### Stable cue representations of CS+ items in TMP and AMY during extinction

Previous fMRI research has shown that the stability of neural representations across repetitions predicts both fear learning (Visser et al., 2013) and episodic memory formation (Lu et al., 2015; Xue et al., 2010). We employed RSA to compare the stability of CS+ vs. CS- cue representations during acquisition and extinction (Figure 3A), again focusing on the time period of cue presentation. We built representational feature vectors based on the pattern of iEEG power values across frequencies (44 values in the 1-100Hz range) and across electrodes in each ROI. These patterns were extracted in windows of 500ms, incrementing in steps of 50ms (90% overlap), and were compared across all pairs of trials using Spearman correlations (see Contrast-based RSA, Methods).

**Figure 3.**
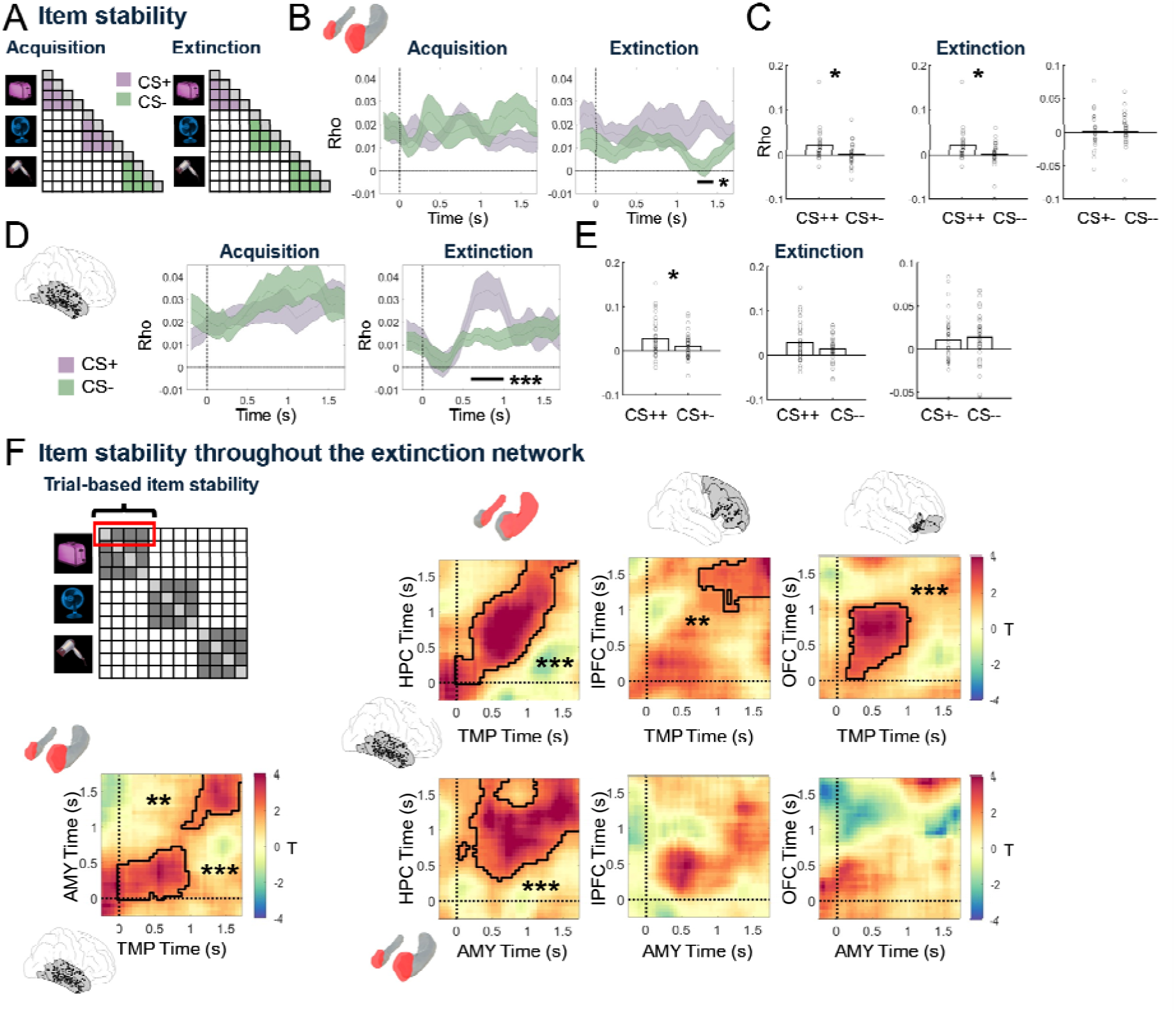
Item stability in lateral temporal cortex, amygdala, and across the extinction network. A) Item stability, defined as the average similarity of neural patterns representing individual items across repeated presentations, was assessed separately for CS+ and CS- trials during acquisition and extinction. B) Analysis of item stability in the AMY revealed no significant differences between CS+ and CS- trials during acquisition (left). During extinction (right), item stability was significantly higher for CS+ as compared to CS- items in a late time period during cue presentation (1.2-1.4s after cue onset). C) Item stability during extinction in the significant time period shown in panel B, right, for CS++ and CS+- (left panel), CS++ and CS-- (middle panel), and CS+- and CS-- trials (right panel). D) Analysis of item stability in the TMP revealed no significant differences between CS+ and CS- trials during acquisition (left). During extinction (right), item stability was significantly higher for CS+ as compared to CS- items in a time period from 0.65s-1s after cue onset. E) Item stability during extinction in the time period shown in panel D, right, for CS++ and CS+- (left), CS++ and CS-- (middle) and CS+- and CS-- (right) trials. F) Top left: A single-trial metric of item stability was computed by assessing the similarity of each trial across repeated presentations (average of the comparisons highlighted with a red rectangle in the RSA matrix). Bottom left: Fluctuations in item stability were coordinated across trials between the TMP and the AMY during the first second of cue presentation and towards the end of the cue presentation period. Right: Item stability was computed in each ROI and correlated across trials with the values observed in TMP (upper row) and AMY (lower row). In panels B and D, black horizontal lines depict significant time periods after correction for multiple comparisons using cluster-based permutation statistics. Time zero indicates the onset of the CS, and shaded lines depict group average Rho values ± S.E.M. Two-sided paired t-tests were applied at every time point (panels B and D) or every time-by-time point (panel F). In panel F, significant regions surviving multiple comparisons corrections using cluster-based permutation statistics are outlined in black. Time zero indicates the onset of the CS in both time axis. * p < 0.05; ** p < 0.01; *** p < 0.001. AMY: Amygdala; HPC: Hippocampus; lPFC: Lateral Prefrontal Cortex; OFC: Orbitofrontal Cortex.

During acquisition, item stability did not differ between CS+ and CS- items in any ROI (all p_corr_ > 0.66). During extinction, we observed significant increases in item stability for CS+ vs. CS- cues in the AMY between 1.25-1.5s (p_corr_ = 0.018), overlapping in time with the period of theta power increases for CS- items (Figure 3B, right). Follow-up analyses within this time period revealed that item stability was significantly higher for CS++ as compared to both CS+- trials (t(31) = 2.30, p = 0.028; Figure 3C, left) and CS-- trials (t(31) = 2.39, p = 0.023; Figure 3C, middle). No differences were observed between CS+- and CS-- trials (t(31) = 1.3, p = 0.2; Figure 3C, right). In the TMP, item stability during extinction was significantly higher for CS+ as compared to CS- trials between 0.65-1s (p_corr_ < 0.005; Figure 3D, right). Within this time window, TMP item stability was significantly higher for CS++ as compared to CS+- trials (t(40) = 2.46, p = 0.018; Figure 3E, left), but no significant differences were observed between CS++ and CS-- trials (t(40) = 1.64, p = 0.11; Figure 3E, middle) or between CS+- and CS-- trials (t(40) = 0.52, p = 0.61; Figure 3E, right). No significant differences in item stability were observed in other ROIs during extinction (all p_corr_ > 0.31).

We next investigated whether item stability was coordinated across brain regions within the extinction network, analogous to the coordination of item-specific representations between HPC and TMP during episodic memory retrieval (Pacheco Estefan et al., 2019). Specifically, we tested whether trial-level metrics of item stability were correlated between AMY and TMP – where condition differences between CS+ and CS- items were observed – and between AMY or TMP and any other ROI. For each trial, ROI, and time point, we averaged the similarity of the representation of a particular item with the similarity of the same item in all other trials (Figure 3F top left, Methods).

Item stability was significantly correlated between TMP and AMY (Figure 3F, bottom left): Trials with high item stability in TMP also showed high item stability in AMY in two temporal clusters, at the beginning of the cue period in both regions (0- 0.95s, p_corr_ < 0.005), and shortly before the presentation of the US (0.9-1.75s, p_corr_ = 0.01). In addition, item stability was coordinated throughout the extinction network (Figure 3F, right): TMP item stability correlated with item stability in HPC, lPFC, and OFC (all p_corr_ < 0.005). AMY item stability correlated with item stability in HPC (p_corr_ = 0.005), but not in prefrontal regions (lPFC: p_corr_ = 0.375; OFC: p = 0.25). Notably, most effects occurred around the diagonal in the temporal generalization map, demonstrating that item stability was coordinated at matching time periods across regions. However, the time windows at which item stability was coordinated varied across regions: While TMP-OFC correlations occurred early after cue onset, TMP- HPC and AMY-HPC correlations occurred across the whole time period of cue presentation, and TMP-lPFC correlations occurred close to the presentation of the US. In supplementary analyses, we tested whether item-stability differed between CS+ and CS- trials, and between CS++ and CS-- trials during extinction, and observed no significant differences between these conditions (Supplementary Note 1). All relationships between all metrics are shown in the summary Figure 8 (see below).

### Context-specific representations in HPC and lateral PFC

Rodent studies have described the crucial role of PFC, in coordination with HPC and AMY, in mediating the context dependency of extinction learning (Gilmartin et al., 2014; Maren et al., 2013; Orsini et al., 2011). We investigated context-specific representations in these regions and across the extinction network, by comparing the similarity of representations of same versus different contexts (Figure 4A, Contrast-based RSA analysis, Methods). We analyzed both the time period when only the context was shown and the subsequent time period when context and cue were shown conjointly, during both acquisition and extinction (Figure 4).

**Figure 4.**
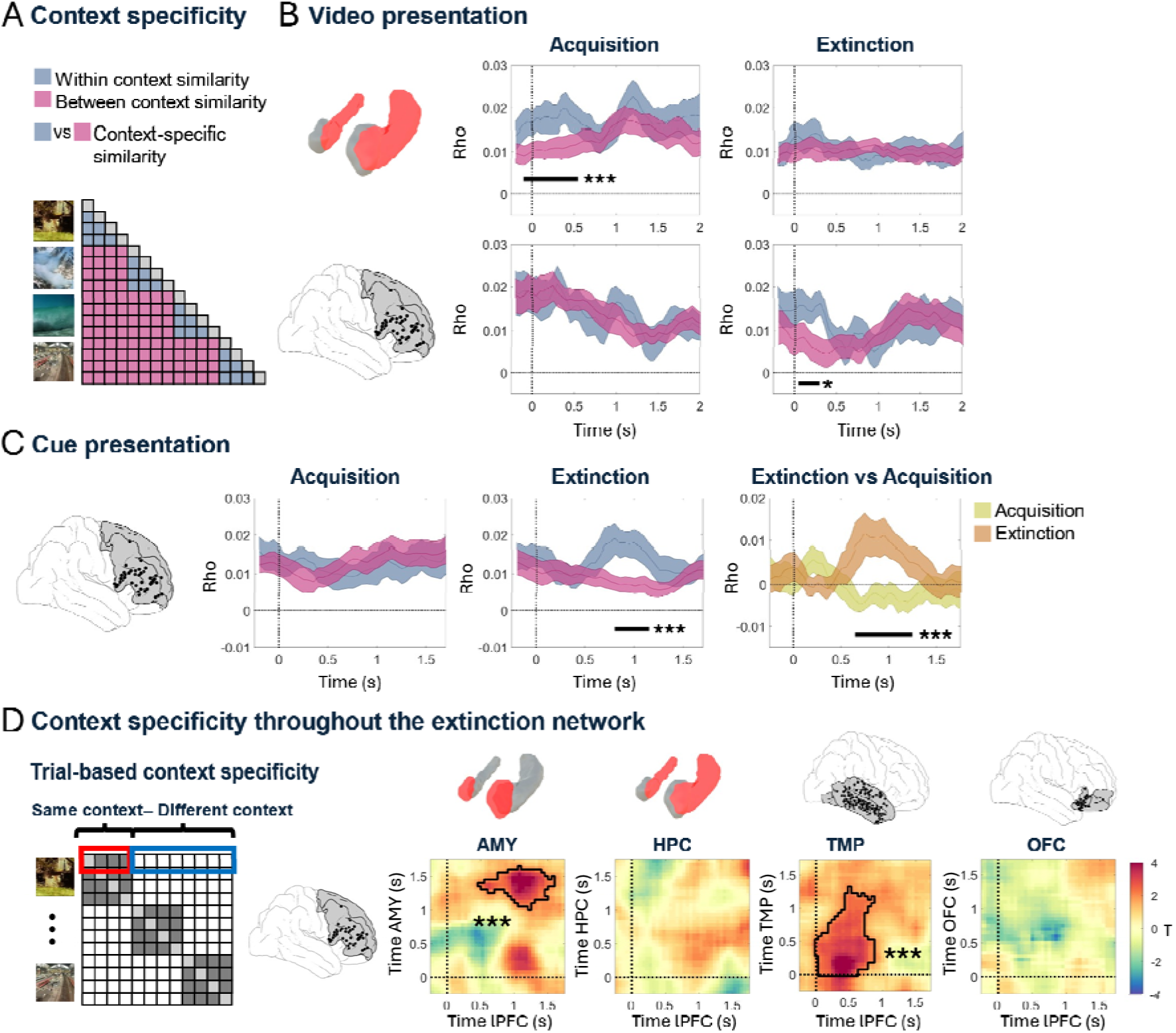
Context specificity in HPC, lPFC and throughout the extinction network. A) Context specificity analysis. Representational patterns corresponding to the presentation of individual context videos were compared across same contexts (blue) and different contexts (pink) during acquisition and extinction. Periods of context specificity were defined as those where same context correlations were significantly higher than different context correlations. B) Context specificity during video presentation in HPC (top) and lPFC (bottom). During acquisition (left), context-specific representations were observed in the HPC locked to the onset of the video. During extinction (right), a similar effect was observed in the lPFC. C) Context specificity during cue presentation. During acquisition (left), no significant differences were observed between same and different contexts. During extinction (middle), context-specific representations were observed in a time period from 0.8 to 1.15s. Right: Context specificity during extinction (brown) was significantly higher than during acquisition (green). D) Left: Trial-specific values of context specificity, defined as average same minus different context correlations in each individual trial, were computed in each of our ROIs. Right: Focusing on the lPFC, we correlated context specificity values between regions and across trials. Significant regions surviving multiple comparisons corrections using cluster-based permutation statistics are outlined in black. In panels B and C, black horizontal lines at the bottom of each panel depict significant time periods after correction for multiple comparisons using cluster-based permutation statistics. Time zero indicates the onset of the CS, and shaded lines depict group average Rho values ± S.E.M. Two-sided paired t-tests were applied at every time point (panels B and C) or every time-by-time point (panel D). AMY: Amygdala; HPC: Hippocampus; lPFC: Lateral Prefrontal Cortex; OFC: Orbitofrontal Cortex.* p < 0.05; ** p < 0.01; *** p < 0.001

During the time period when only the context was shown, we observed significant context-specific representations in the HPC during acquisition (0-500ms; p_corr_ < 0.005; Figure 4B, top left) and in the lPFC during extinction (50-300ms, p_corr_ = 0.0245; Figure 4B, bottom right). No significant context-specific representations were observed in the other ROIs during acquisition or extinction (all p_corr_ > 0.06).

During the subsequent period when both contexts and cues were shown, we did not observe any significant context-specific representations during acquisition in any ROI (all p > 0.125). During extinction, however, we observed significant context-specific representations in lPFC (0.8s-1.15s; p_corr_ = 0.012), partially overlapping with the time period showing item stability in TMP (Figure 4C, middle). lPFC context specificity was significantly higher during extinction than acquisition (p = 0.001; Figure 4C, right). No other ROI showed context specificity in this time window during extinction (all p > 0.23).

To investigate whether context specificity was coordinated across the extinction network, we again performed a trial-level correlation analysis between the HPC, the lPFC, and other ROIs, consistent with our analysis of item stability coordination (Figure 3). Specifically, we computed a trial-level metric of context specificity by calculating the average difference in correlations between a given trial and other trials with the same context vs. other trials with a different context (Figure 4D, left; Methods). We focused on the regions and time periods where we observed significant context-specific effects in our previous analysis: the HPC when only the video was shown during acquisition, and the lPFC when both video and cue were shown during extinction. We correlated the values observed in these two regions with those found in our other ROIs.

Context specificity in the HPC was not coordinated with any other ROI (all p_corr_ > 0.06). During extinction, we observed a significant coordination of context specificity in lPFC and TMP during an early time period (0-1.3s) and between lPFC and AMY during a late time period (0.5-1.55s; all p_corr_ < 0.005; Figure 4D, rigth). Both periods overlapped with the time period showing significant context specificity in lPFC, and the time period of lPFC-AMY coordination also overlapped with the AMY theta power effect. Correlations between lPFC and other ROIs were not significant (all p > 0.24). We did not observe significant differences between conditions in the coordination analysis of context-specificity (Supplementary Note 1).

Taken together, we found significant context-specific representations during both acquisition and extinction, but in different brain regions (HPC during acquisition, lPFC during extinction) and at different time periods (during acquisition: when the context was presented alone; during extinction: when the context was shown with the cue). During extinction, the magnitude of context specificity was coordinated across the extinction network between the lPFC and regions showing theta power differences (AMY) and/or differential item stability between CS+ and CS- (AMY and TMP). In a next step, we evaluated whether the different representational metrics were related.

### Links between amygdala theta power, item stability in AMY and HPC, and context specificity in lPFC

Our results so far indicate that, during the extinction phase, both sensory TMP regions and AMY represent CS+ cues in a more stable way than CS- cues. They also show that the lPFC exhibits context-specific representations in an overlapping time period. In order to uncover the relationship of AMY theta power (AMY_THETA_), TMP/AMY item stability (TMP_ITEM_ and AMY_ITEM_), and lPFC context specificity (lPFC_CONTEXT_), we investigated their correlation across trials, and extended this analysis to item stability and context specificity effects in other ROIs during extinction (Figure 5A). We extracted a single-trial metric of AMY_THETA_ based on the cluster of significant differences between CS+ and CS- items observed during extinction (Figure 2; see as schematic depiction in Figure 5A, left). We separately z-scored this metric for CS+ and CS- trials, in order to avoid any spurious correlation of single-trial values driven by main condition differences. We first focused on the time period of significant AMY theta power effects and averaged item stability and context specificity across this time period in each trial and each ROI.

**Figure 5.**
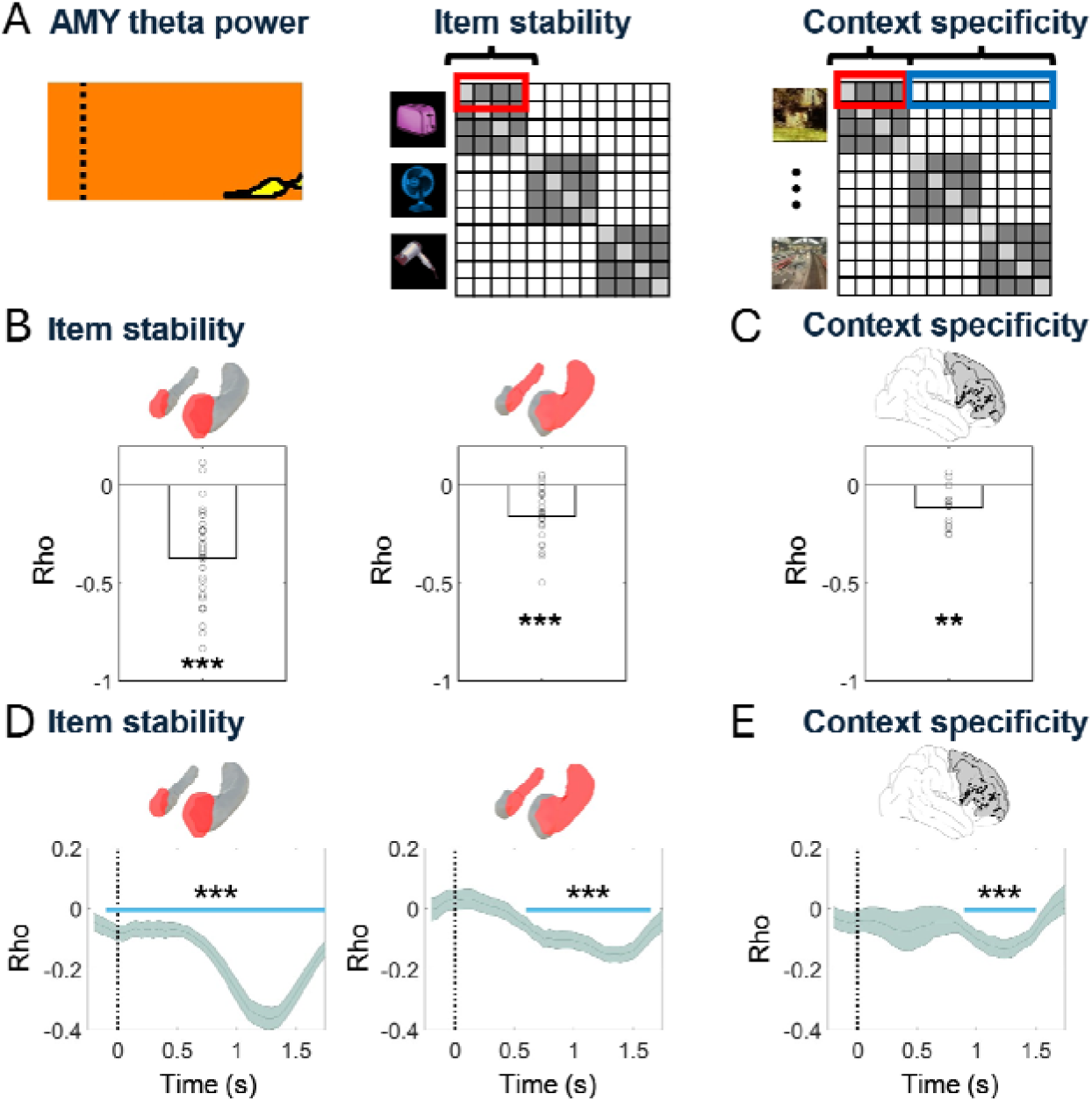
Amygdala theta power correlates with item stability and context specificity across the extinction network. A) Top left: Amygdala theta power was averaged in the time frequency cluster where differences between CS+ and CS- items were observed in the power analysis during extinction. Left: schematic depiction of the selected time frequency period (highlighted in yellow). The average theta power in this cluster was correlated across trials with two metrics: item stability (middle), and context specificity (right). Group level Rho values were contrasted against zero using paired t-tests. B) AMY theta power was correlated across trials with item stability in both AMY (left) and HPC (right). C) AMY theta power also correlated with lPFC context specificity across trials. D) Time-resolved analysis. Left: AMY theta power and item stability correlated throughout the entire cue presentation period. Right: Correlations between AMY theta power and HPC item stability were observed during a later time window (600ms to 1.65s) of cue presentation. E) AMY theta power correlated with lPFC context specificity during a time period that overlapped with the significant condition differences in theta power in AMY (900ms-1.5s). In panels D and E, shaded lines depict group average Rho values ± S.E.M, and time zero marks the onset of cue presentation during the extinction phase. ** p < 0.01; *** p < 0.001

We observed that across trials, AMY_THETA_ correlated with both AMY_ITEM_ (t(31)= -9.08; p_corr_ = 1.54e-9; Figure 5B, left) and HPC_ITEM_ (t(23) = -5.61; p_corr_ = 5.25e-5; Figure 5B, right), but not with item stability in any other ROI (all t < 0.79; all p > 0.44). Notably, AMY_THETA_ also correlated with lPFC_CONTEXT_ (t(14) = -4.37; p_corr_ = 0.0032; Figure 5C). Context specificity in none of the other ROIs correlated with AMY_THETA_ (all t < 1.43, all p > 0.16). To assess the temporal specificity of these effects, we performed the same analysis across the entire cue presentation period, focusing on the ROIs where we observed significant effects (i.e., AMY and HPC for item stability analysis and lPFC for context specificity). Notably, we observed significant correlations during the whole time period of cue presentation in AMY, which were most pronounced when the AMY_THETA_ condition difference effects were observed (i.e., from 1.18-1.75s after cue presentation; p_corr_ < 0.005; Figure 5D, left). Correlations between AMY_THETA_ and HPC_ITEM_ reached significance from 600ms to 1.65s (p_corr_ < 0.005; Figure 5D, right), and correlations of AMY_THETA_ with lPFC_CONTEXT_ were observed in an intersecting time period (from 900ms to 1.5s after cue onset; p_corr_ < 0.005; Figure 5E). In supplementary analyses, we tested whether correlations of theta power and item-stability or context specificity differed between CS+ and CS- trials, and between CS++ and CS-- trials during extinction, focusing on the regions where we observed significant effects in the analysis including all trials. Our results revealed no statistical differences between conditions (Supplementary Note 2). Taken together, these results demonstrate that AMY_THETA_ was correlated with both AMY and HPC item stability and lPFC context specificity during a late time period of cue presentation.

### LPFC context specificity predicts reinstatement of fear memory traces in TMP

Both rodent studies (Gilmartin et al., 2014; Maren et al., 2013) and clinical studies in humans (Craske et al., 2014; Garfinkel et al., 2014; Milad & Quirk, 2012; Wang et al., 2024) suggest that high context specificity during extinction predicts renewal, i.e., a reoccurrence of fear memories in acquisition contexts or novel contexts. We thus quantified the degree to which the representations of the three cue types during acquisition and extinction re-appeared during the test phase. Specifically, we computed the similarity of every trial during acquisition (or extinction) with all trials in which the same item was shown during the test phase. Averaging these values across trials separately for each cue type (CS++; CS+-; CS--) and subtracting acquisition reinstatement and extinction reinstatement yielded a subject-specific metric of fear reinstatement (REINST = REINST_ACQ_ – REINST_EXT_; Figure 6A). We correlated these reinstatement values in AMY and TMP, where the main effects of item stability were observed, with subject-averaged metrics of context specificity in lPFC during the extinction phase (Figure 6B).

**Figure 6.**
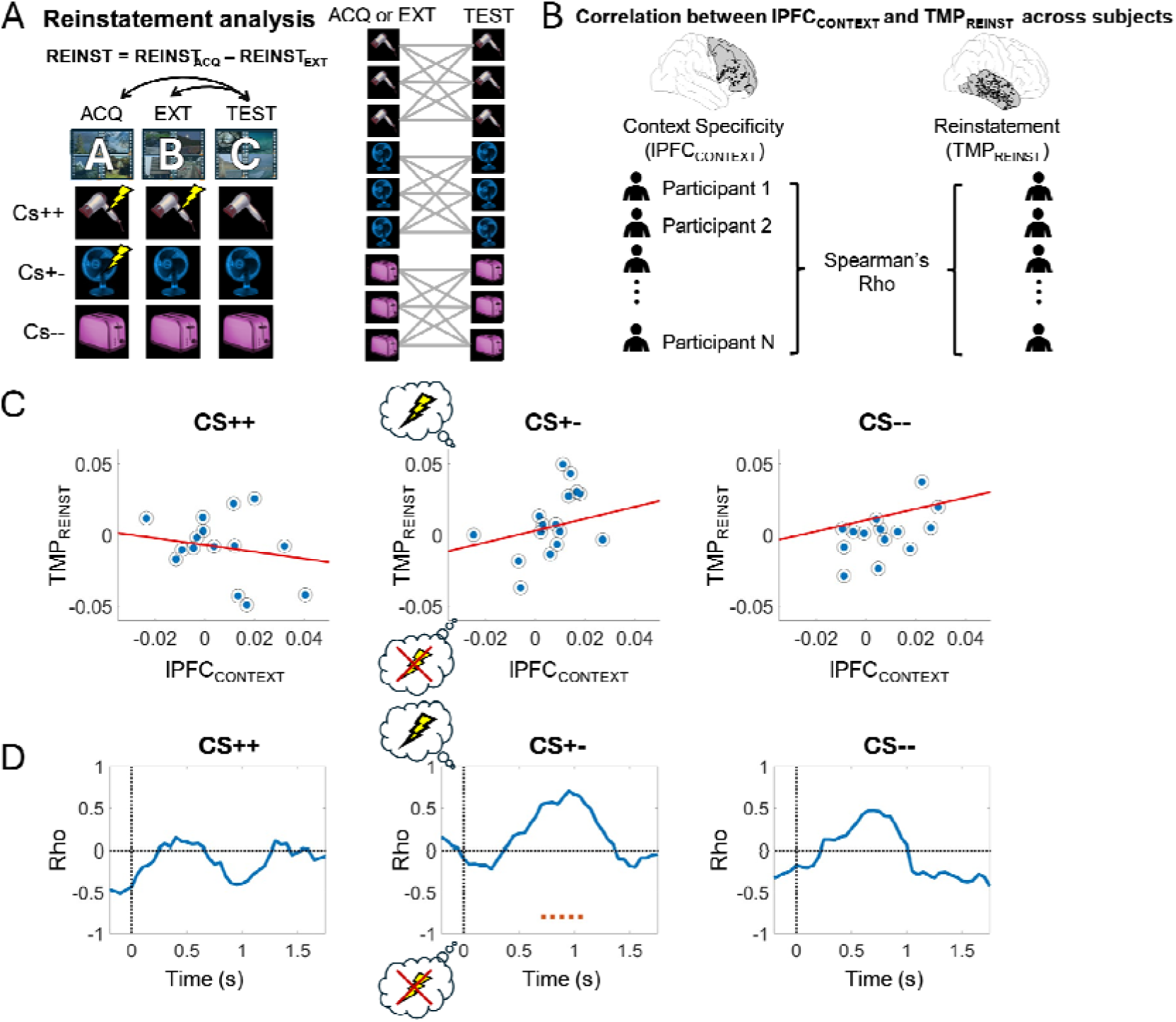
Context specificity shifts the balance in reinstatement of fear vs. extinction memory traces. A) Reinstatement analysis. Reinstatement of acquisition and extinction activity patterns during the test was calculated separately for CS++, CS+- and CS-- items (left), by computing the similarity across repeated exposures to the same item across experimental phases (right). Acquisition and extinction reinstatement values were subtracted and averaged across trials, resulting in a single value of differential reinstatement in each subject and condition (REINST = REINST_ACQ_ – REINST_EXT_). B) Context specificity in lPFC and reinstatement in TMP were correlated across our group of participants using Spearman correlations. C) LPFC context specificity and TMP reinstatement were correlated during the time period of significant context-specific effects in lPFC. Note that for CS+- trials, positive values of TMP reinstatement indicate predominant reinstatement of fear memory traces, while negative values indicate predominant reinstatement of extinction memory traces. D) Context specificity in lPFC was correlated with TMP reinstatement using overlapping 500ms windows with a 95% overlap (i.e., shifted by 50ms). A cluster of significant correlations was identified specifically CS+- trials (middle panel), which coincided with the time period of lPFC context-specific effects. Orange dashed line in panel D, middle, indicates significant regions surviving multiple comparisons correction at p_corr_ = 0.052.

First, we analyzed possible relationships between lPFC context specificity and reinstatement in AMY or TMP during the time period when significant context-specific effects were observed during extinction. As hypothesized, we observed a positive correlation between context specificity in lPFC and reinstatement in TMP, indicating more pronounced reinstatement of acquisition than extinction memory traces in participants with more pronounced context specificity during extinction. Interestingly, this effect occurred specifically for CS+- trials whose contingencies changed from acquisition to extinction (Rho(15): 0.57, p = 0.024; Figure 6C, middle), but not for CS++ (Rho(14) = -0.13, p = 0.3; Figure 6C, left) or CS-- trials (Rho(15) = 0.4, p = 0.13; Figure 6C, right). LPFC context specificity did not correlate with AMY reinstatement for any cue type (CS++: Rho(13) = -0.01 p = 0.98; CS+-: Rho(13) = 0.029, p = 0.92; CS--: Rho(13) = 0.28, p = 0.32). This effect was not observed when we correlated lPFC context specificity with either TMP REINST_ACQ_ or REINST_EXT_ separately for any of our three trial types (Acquisition: all p > 0.229; Extinction: all p > 0.56), suggesting that it specifically affects the balance (or competition) between reinstatement of acquisition vs. extinction memory traces (see Discussion).

Next, we extended the analysis of reinstatement to all time points per trial for our three trial types (Figure 6D). We correlated lPFC context specificity with averaged levels of TMP reinstatement in windows of 400ms (same size as the lPFC context specificity effect), incrementing in steps of 50ms, and correlated these two metrics across subjects for each time window. We corrected for multiple comparisons using cluster-based permutation statistics, by shuffling the subject id labels (Methods). We observed a very specific window where increases in correlations were observed at trend level and specifically in CS+- trials, from 700ms to 1.05s (p = 0.055; Figure 6D, middle). These results suggest that the relationship between lPFC context specificity and TMP reinstatement in CS+- trials occurred selectively during the time periods of significant context specificity in lPFC.

In complementary analyses, we evaluated whether overall levels of acquisition-to-test, extinction-to-test and differential (acquisition-to-test minus extinction-to-test) reinstatement differed across trials, and whether for each cue type, acquisition-to-test and extinction-to-test reinstatement differed in the TMP and the AMY. We observed a trend for higher acquisition-to-test reinstatement than extinction-to-test reinstatement in the AMY specifically in the CS+- trials, suggesting that fear memories of CS+- items might show slightly higher levels of reinstatement than the subsequent extinction memories of these items (Supplementary Note 3 and Supplementary Figure 1).

### Reinstatement of extinction memory traces in TMP predicts safety responses

In our final analysis, we evaluated whether reinstatement of extinction memory traces during the test phase predicted subjective ratings of safety. For every item during the test phase, similarity was calculated with all instances of this item during extinction, and these values were correlated across trials with the ratings during the test phase. We found that reinstatement of extinction memory traces was positively correlated with subjective safety ratings for CS+- cues (t(18)= 2.3; p = 0.0078), while this was not the case for CS++ (t(20)= 0.35; p = 0.72) or CS-- cues (t(22)= -0.16; p = 0.87).

## Discussion

Here we investigated the neurophysiological mechanisms and representational signatures of extinction learning in the human brain. CS- items during extinction were associated with increases in theta power in the AMY and lower levels of item stability in the TMP. Moreover, extinction learning showed higher context specificity in lPFC, which predicted the reinstatement of fear memory traces in TMP. Further analyses unraveled the distinct pattern and time-course of coordinated representations across the fear and extinction network, which is summarized in Figure 8.

### A safety signal during extinction in the AMY correlating with representational changes of contexts and items

While a previous human iEEG study reported increases in theta power for CS+ vs. CS- items during acquisition (Chen et al., 2021), our results showed higher theta power for CS- vs. CS+ items during extinction. This apparent discrepancy may be explained by the differing cognitive demands of acquisition and extinction, with the latter requiring pronounced coordination between AMY and other brain regions coding for contexts and cognitive control such as HPC and PFC. Indeed, increases of AMY theta power during extinction were correlated with higher levels of item stability in both AMY and HPC and with higher levels of context specificity in lPFC. The lack of AMY theta power effects during acquisition in our study contrasts with findings in animals and humans, which have typically reported theta increases during the formation of cue-US associations across several regions of the fear and extinction network (e.g., Chen et al., 2021; Pirazzini et al., 2023). Several factors may explain this discrepancy, including differences in recording methodologies (iEEG versus EEG), the distinct regions targeted (e.g., different subregions of the PFC targeted in [Chen et al., 2021]), interspecies differences in theta oscillations (Jacobs, 2014), and the use of a more “cognitive” paradigm in our study with a milder aversive stimulus.

In our paradigm, both CS+- and CS-- trials signal safety during extinction – either newly learned safety following extinction (for CS+- trials) or consistent safety throughout the experiment (for CS-- trials). Thus, the successful extinction of previous fear should result in neural activity and representations of CS+- items that match those of CS-- items—in line with our findings on both AMY theta power and AMY and TMP item stability.

Notably, the increase in theta power for CS- items observed during extinction, coupled with the absence of such increases during acquisition, suggests that theta power does not simply signal safety but rather safety within the specific context of extinction. At least two key differences distinguish acquisition from extinction in our paradigm: (1) Contingencies changed for one of the cues (CS+-), and (2) contexts changed from A to B. While both of these factors may have influenced theta responses, future research is needed to determine whether theta power during extinction signals safety specifically when both context and contingencies change in the context of a multi-cue paradigm such as ours, or whether it would also be observed in extinction learning paradigms in which contexts do not change (e.g., AAB designs) or in those including only one cue (CS+-).

We note that in our AMY theta power analyses, some time-frequency bins observed early in the trial (∼300-500ms) showed numerical increases in theta power for CS+ items at an uncorrected level. This might seem surprising, as it contrasts with the main effect observed later in the trial (1.18-1.75s) which reflects a decrease in theta power for threatening items. However, these numerical increases are within the range of what is expected by chance, as assessed by our cluster-based permutation analysis. While our primary analyses focused on theta frequency oscillations, humans are able to rapidly distinguish and respond to threats and safety cues (Méndez-Bértolo et al., 2016). This raises the possibility of a potential role of high-frequency neural activity in signaling threat and safety. Indeed, theoretical accounts suggest that fast cycle gamma oscillations might promote fast information transfer between brain regions (Fell & Axmacher, 2011), facilitating fear learning by modulating the timing of neuronal firing inducing plasticity (Cattani et al., 2024). In particular, gamma cycles align with several biophysical factors regulating excitatory input integration in basolateral amygdala principal neurons (Bocchio et al., 2017), and experimental studies in rodents have shown that amygdala gamma oscillations contribute to fear memory formation through theta-gamma coupling and spike-field coherence (Popescu et al., 2009; Stujenske et al., 2014). While we did not observe any effect of gamma frequency oscillations in our paradigm, we believe further research will be needed to fully assess their possibly role in signaling threat and safety and modulating fear learning in humans—for example, via phase amplitude coupling (Saint Amour di Chanaz et al., 2023). Interestingly, recent findings suggest that hippocampal representational patterns linked to amygdala gamma bursts reinstate the content of emotional memories (Costa et al., 2024), providing evidence for a potential role in the representation of fear in the interaction with the hippocampus

The high temporal resolution of iEEG recordings allowed us to shed light on the relative time courses of signals across regions of the extinction network. Our results reveal a temporal sequence in which an earlier representation of specific contexts in lPFC is followed by a later correlation of lPFC_CONTEXT_ and AMY_CONTEXT_ across trials, which, in turn, coincides with the safety-related theta power increase in AMY. This temporal order of representational and neurophysiological signals aligns with previous research showing that distributed areas of the extinction network, including the PFC, first coordinate the processing of contextual information, which subsequently influence the AMY (Holland & Bouton, 1999; Quinn et al., 2008).

Notably, the correlation of lPFC and AMY context-specificity overlapped with the period of lPFC context specificity (from 0.8 to 1.15s after cue onset). However, it also shows a delay between lPFC and AMY, with lPFC context specificity emerging ∼1s earlier. This temporal lag seems to decrease during later trial periods. The main effect of lPFC context-specificity occurs immediately prior to the time period when the strongest coordination is observed with the AMY (at approximately 1.2s). Although the AMY did not show a main effect of context specificity—possibly because the similarity of same-context representations was attenuated by repetition suppression—the observed temporal offset suggests that the effect observed in the lPFC drives the AMY representations. Our findings underscore the relevance of dynamic and distributed context representations throughout the extinction network during fear extinction. More specifically, they suggest that AMY safety signals and representations during extinction depend on the preceding representation of specific contexts in lPFC.

### The context-dependency of extinction learning shifts the balance in the competition between fear and extinction memory traces towards a renewal of fear

Our findings also shed light on the functional role of lPFC context representations in modulating the reinstatement of fear vs. extinction memories. Reinstatement of memory traces has previously been analyzed via encoding-retrieval similarity (Pacheco Estefan et al., 2019; Staresina et al., 2016; Yaffe et al., 2014). In the context of fear extinction, previous fMRI studies investigated the reinstatement of fear and extinction memory traces using multivoxel pattern similarity analysis (Hennings et al., 2022). Following a similar approach, here we implemented a novel metric of reinstatement that directly compared the reoccurrence of fear vs. extinction memory traces, assuming an inhibition of these two memories during test (Bouton & Swartzentruber, 1991; Lebois et al., 2019; Santini et al., 2008; Szeska et al., 2020). Using this metric, we observed that higher fear vs. extinction memory reinstatement in TMP correlated with context-specific representations in the lPFC – i.e., in participants with elevated levels of context-signaling in the lPFC during extinction, the balance between fear memory reinstatement and extinction memory reinstatement was shifted towards the former (Figure 6). Importantly, this effect was selectively observed for CS+- cues, but not for CS++ or CS-- cues, which are unlikely to compete between acquisition and extinction. Our finding aligns with clinical observations that high levels of context dependency during extinction reduce the persistence of the safety responses learned during extinction, and favor the recurrence of fear (Lebois et al., 2019). One may assume that if extinction contexts are represented more specifically, the newly formed extinction memories are considered as an exception, and the original fear memory is likely to come back. Our results further show that aversive memories are extinguished through an interaction between executive control (lPFC) and sensory regions (TMP) representing contexts and items, respectively. Possibly, more pronounced context representations in lPFC involve effortful inhibition processes similar to those during episodic memory control (Engen & Anderson, 2018), which are difficult to maintain in the longer run.

### Fear learning, extinction, and episodic memory

Our results highlight the crucial role of both HPC and lPFC in encoding contextual information during fear learning and extinction, consistent with findings in rodents (Bouton, 2004; Gilmartin et al., 2014; Maren et al., 2013). Notably, the lPFC was selectively engaged in the representation of context-specific activity during extinction and not during acquisition, again consistent with its involvement in the suppression of fear memories via context-signaling. While we observed a context-specific representation during acquisition as well (in the HPC), this effect did not coincide with the time period of the cue presentation – suggesting that specific contexts are not neglected during acquisition but exert a less prominent influence on the memory traces of individual cues. While the lPFC effects align with the role of this region in inhibiting fear expression (Gilmartin et al., 2014; Maren et al., 2013), the HPC responses are consistent with the greater sensitivity of this region to views of landscapes and scenes (Maguire & Mullally, 2013; Rolls, 1999) and the well acknowledged role of the HPC in rapidly encoding new memory traces during acquisition (McClelland et al., 1995). We note that several studies have shown that fear acquisition is characterized by an overgeneralized and decontextualized fear responses, while extinction engages PFC-dependent mechanisms more reliant on context representation (for reviews, see Gilmartin et al., 2014; Maren et al., 2013). We previously found that overgeneralization of representations predicted subsequent intrusive memories (Kobelt et al., 2024), which are notorious for their decontextualized nature and the fact that they can be easily and involuntarily triggered by ubiquitous sensory cues.

Our findings suggest that extinction memory traces involve neurophysiological patterns and representational characteristics that differ from those formed during acquisition and are more reminiscent of episodic memory traces (see also Hennings et al., 2022). First, CS- cues were associated with increased theta oscillations during extinction. Previous studies on the role of theta oscillations for episodic memory formation suggest that theta power increases particularly for associative – i.e., context-dependent – memories, while the encoding of item memories is commonly associated with theta power reductions (Herweg et al., 2020). Second, the increased levels of lPFC context-specificity during extinction are reminiscent of the formation of new episodic memories, which are defined, among other factors, by their dependence on context (Tulving, 2002; Yonelinas et al., 2019). These findings support the notion of fear acquisition as a form of associative learning that lacks contextual specificity, while fear extinction, similar to episodic memory, is more strongly influenced by contextual representations. Third, memory traces of CS- items during extinction were less stable and, therefore, more trial-specific in sensory regions of the TMP, reminiscent of the unique and distinct representations during individual trials observed in episodic memory paradigms. Indeed, while previous studies have linked the stability of neural representations to both episodic memory encoding (Lu et al., 2015; Xue et al., 2010) and the formation of fear associations (Visser et al., 2013), others have suggested that item stability is modulated by contextual factors and is disrupted when incongruent contextual information is presented (Wu et al., 2023). To summarize, these findings point towards the flexible and malleable nature of safety memory traces during extinction; and indeed, higher level of context-dependency (and thus putatively higher “episodicity” of extinction memories) predicted their relative weakening as compared to the more robust and generalized initial fear memory traces.

While both extinction learning and episodic memory are tightly linked to contextual information, these two forms of associative learning certainly differ in other respects. Episodic memory is specifically defined by its connection to “mental time travel” – the ability to mentally project oneself into the past or future and to reconstruct specific experiences (Tulving, 2002) – a feature that is not evident in fear extinction paradigms. Moreover, fear learning and extinction typically occur across repeated trials, while episodic memory has been characterized as one-shot learning (McClelland et al., 1995). Despite these differences, our data shows that extinction learning shares representational signatures with item-context associations in episodic memory (Eichenbaum, 2017b; Pacheco Estefan et al., 2019).

### Methodological considerations

Cognitive impairments in implanted epilepsy patients are generally a concern when conducting cognitive experiments using iEEG data. To facilitate the interpretation of our findings, we provide in Supplementary Table 1 all available demographic and clinical information of our patients, including global intelligence quotient (IQ), memory quotient (MQ; assessed via the Wechsler Memory Scale; WMS-III) and seizure onset zone (SOZ). While the IQ scores were within the normal range, the patients in our study showed a lower-than-average MQ. Although our paradigm did not directly probe the specific memory functions assessed by the WMS, this limitation is inherent to intracranial EEG recordings in epilepsy patients and should be acknowledged. While we cannot entirely rule out the influence of memory impairments on our results, the simplicity of our task design—with only three cues and a single instruction for all trials—, and the fact that learning curves align with expected behavioral patterns partially alleviate this concern. Additionally, fear associations can be formed in the absence of conscious awareness of the CS-US contingencies (Knight et al., 2003, 2009), suggesting that explicit memory impairments may not necessarily prevent the formation and extinction of fear associations. Indeed, our trial-by-trial progressive learning analysis confirms that patients successfully understood the task instructions and appropriately learned the CS-US contingencies, adaptatively adjusting their responses to contingency changes in CS+- trials (see Figure 1D).

We note that, similar to other fear conditioning paradigms conducted with humans (e.g., Balooch et al., 2012; Bandarian Balooch & Neumann, 2011; Kinner et al., 2016; Lissek et al., 2008; Lissek & Tegenthoff, 2024; Neumann et al., 2007; Neumann & Kitlertsirivatana, 2010; Shiban et al., 2013), we chose to rely on self-reported ratings of cue ‘threat’ and ‘safety’ as our behavioral measure of fear rather than employing implicit physiological recordings. While explicit ratings have limitations, we decided to employ them based on both ethical and clinical considerations, particularly given the constraints of conducting research with implanted epilepsy patients. Specifically, the use of a mild aversive stimulus (a scream) rather than electric shocks in our study precluded the use of some traditional implicit measures, such as skin conductance or heart rate responses, which are more suited for capturing stronger autonomic reactions. Importantly, research indicates that explicit ratings are more stable than physiological responses across repeated measurements (Corneille & Gawronski, 2024), and several reviews support their validity in fear conditioning research (Boddez et al., 2013; Constantinou et al., 2021). Explicit self-reports have been shown to correspond with both fear startle (Balooch et al., 2012; Lissek et al., 2008) and skin conductance responses (Constantinou et al., 2021; Shiban et al., 2013; Taschereau-Dumouchel et al., 2020). Moreover, self-reports might better capture deliberative processes than implicit measures (Lonsdorf et al., 2017), which aligns with the more “cognitive” nature of our paradigm (see also Kinner et al., 2016; Lissek & Tegenthoff, 2024). Although explicit ratings are susceptible to demand effects or participant interpretation, such effects are unlikely to have played a major role in our results due to the simplicity of the task and the absence of explicit experimenter expectations beyond learning the correct CS-US associations. Crucially, participants were instructed to rate the ‘safety’ of the cue itself rather than their internal emotional state, minimizing subjective bias. Furthermore, apart from analyzing the correlation of the ratings with reinstatement in the TMP (Figure 7), we did not rely on the ratings to categorize stimuli or correlate them with the neural data in other analyses, which minimizes their influence in the overall interpretation of our findings. Finally, we highlight that the trial-by-trial learning effects we observed based on the ratings align closely with what would be expected for skin conductance responses (see for example, Ma et al., 2025).

**Figure 7.**
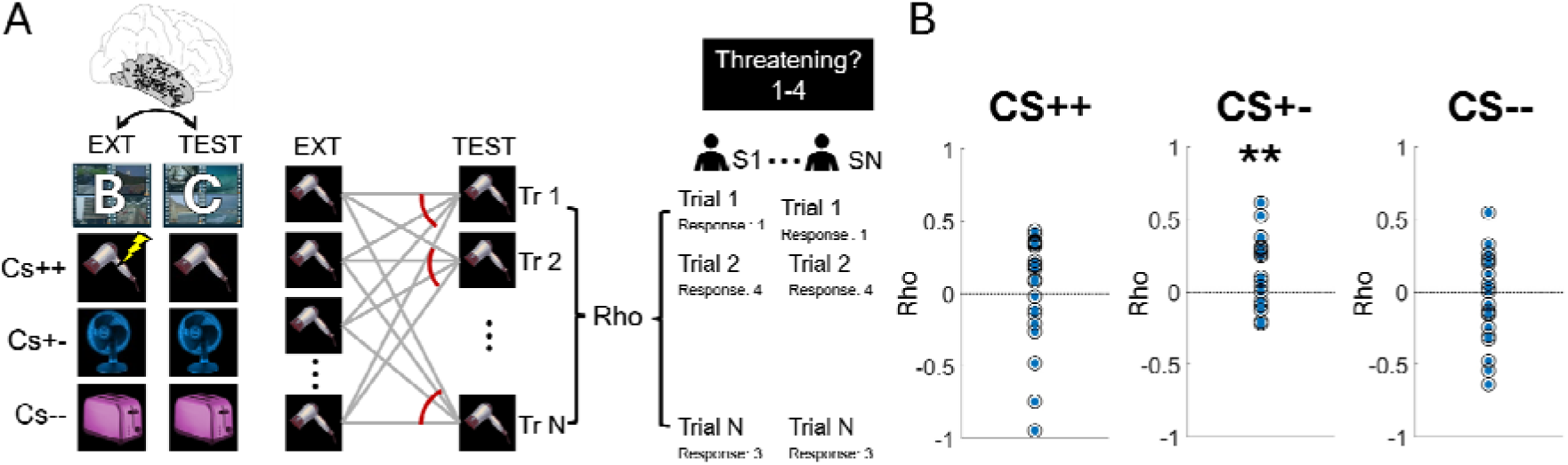
Reinstatement of extinction memory traces in TMP predicts subjective safety ratings. A) Extinction-to-test reinstatement analysis: For each trial type (CS++, CS+-, and CS--), the similarity between extinction and test trials was computed in TMP (left). This was done by averaging the similarity of one trial during the test across repeated presentations of this same item during the extinction phase (middle). Spearman correlations were then computed between extinction-to-test similarity and subjective ratings during the test across trials, in each subject independently (right). B) Fisher Z-transformed correlations were analyzed at the group level to determine whether they significantly differed from zero, separately for CS++ (left), CS+- (middle), and CS-- (right) trials. ** p < 0.01.

**Figure 8:**
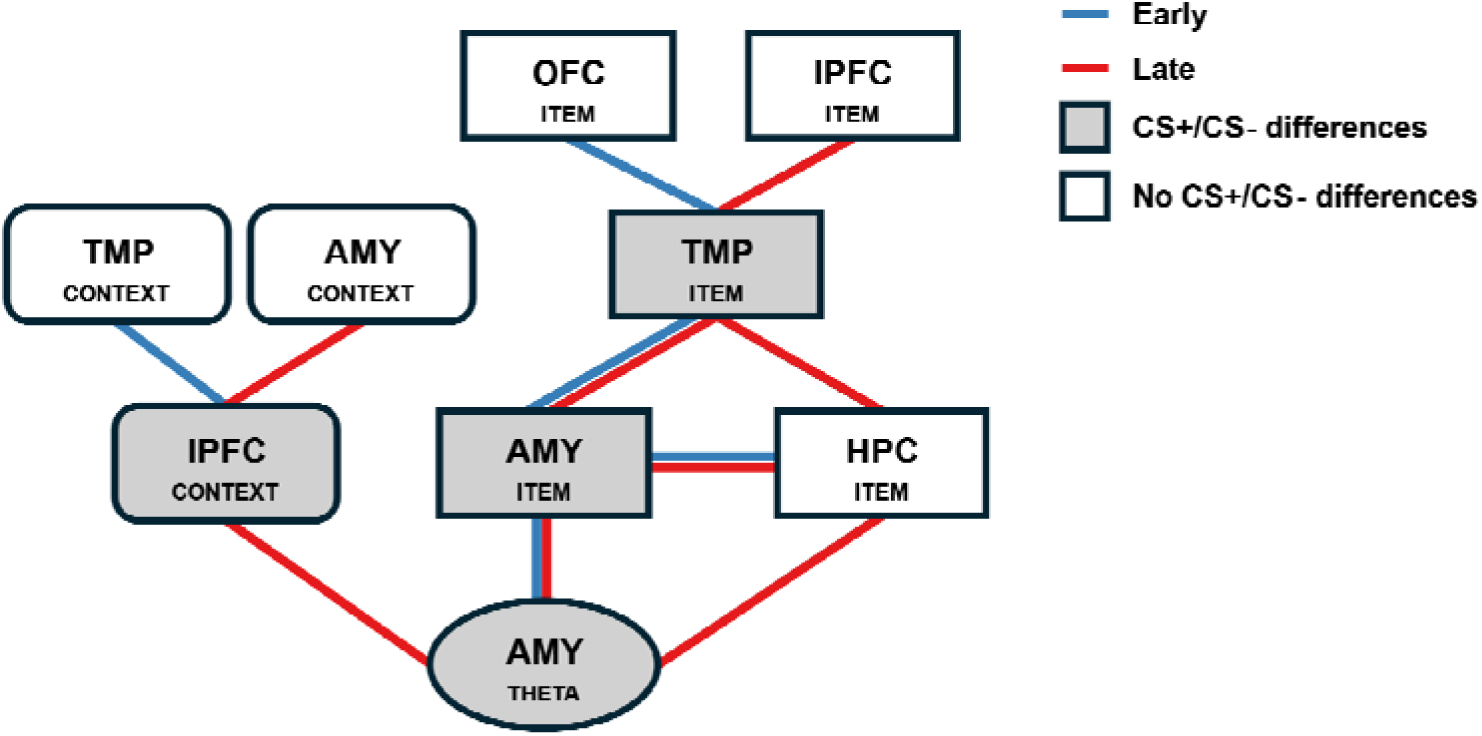
Summary of the main results during extinction learning. The figure summarizes the main results obtained in the power analysis, the item stability analysis and the context specificity analysis during the extinction period across our five ROIs. During the time period of theta power condition differences in the amygdala (CS+ vs CS-; labeled ‘AMY_THETA_’ in the figure), we also observed lPFC context-specific effects (‘lPFC_CONTEXT_’) and item stability effects in AMY and lPFC (‘AMY_ITEM_’; ‘lPFC_ITEM_’). Note that AMY_THETA_ effects occurred during a late time period (indicated with red lines in the figure), but AMY_THETA_ and AMY_ITEM_ also correlated earlier in the trial (indicated with blue lines in the figure). In turn, AMY_ITEM_, TMP_ITEM_ and HPC_ITEM_ effects were observed during early and late time periods. AMY_CONTEXT_ correlated with lPFC_CONTEXT_ during a late time period only, but lPFC_CONTEXT_ correlated with TMP_CONTEXT_ during an earlier time period. Metrics showing main condition differences between CS+ and CS- items during extinction are highlighted in grey. AMY: Amygdala; HPC: Hippocampus; lPFC: Lateral Prefrontal Cortex; OFC: Orbitofrontal Cortex.

Our study was not preregistered, and we acknowledge this as a limitation, as preregistration enhances the transparency and reproducibility of research findings. However, the exploratory nature of our study, which is the first to investigate context-dependent extinction learning in humans via iEEG, made preregistration challenging. Our research aimed to provide a first characterization of the representational dynamics underlying the context-dependent nature of extinction learning—which required the integration of methodologies from diverse domains and the development of novel metrics. For example, we adapted RSA, an established method in episodic memory research, to examine context specificity and item stability within a classical ABC context-dependent extinction paradigm (Figure 2 and 3). We also developed a new measure to quantify fear reinstatement in the context of classical conditioning (see Figure 6). Despite the pioneering and exploratory nature of our study, we note that we formulated specific predictions in the introduction and results sections.

Notably, some of our results were not in line with our predictions. Specifically, we anticipated to observe increased theta power for CS+ as compared to CS- items during acquisition in AMY, which we did not find. Instead, we observed a relative decrease of power for these trials during extinction in this region (see section above: *A safety signal in the AMY driven by context representations in the extinction network*). We also expected context-specific signals both in lPFC and HPC during extinction based on previous literature (Maren et al. 2013). However, context specificity was only observed in the lPFC during extinction, while the involvement of the HPC was only observed during acquisition (see section *Fear learning, extinction, and episodic memory* above). Finally, we expected lPFC context specificity to drive context representations in AMY and HPC, but while lPFC and AMY context representations were indeed correlated across trials, we did not find a main effect of context specificity during extinction in either AMY or HPC (see section *A safety signal during extinction in the AMY correlating with representational changes of contexts and items above*). Despite its exploratory nature, our work represents an important first step in exploring the representations underlying context-dependent extinction learning in humans using iEEG, providing a foundation for more targeted, hypothesis-driven research in the future. In summary, our findings shed light on the key neurophysiological mechanisms, representational characteristics, and functional relevance underlying context-dependence fear extinction in the human brain. They bridge the gap between animal and human studies, contribute first evidence to a novel conceptual framework that integrates fear conditioning and episodic memory research, and provide a mechanistic basis for understanding the renewal of fear in clinical contexts.

## Methods

### Participants

Forty-nine patients (22 females, 28.2 ± 8.18 years) with medically intractable epilepsy participated in the study. Data were collected at the Pitié Salpêtrière hospital, Paris, and the South China Normal University Hospital, Guangzhou, China. The study was conducted according to the latest version of the Declaration of Helsinki and approved by the institutional review board at the Pitié Salpêtrière hospital (C11-16, C19-55 National Institute of Health and Medical Research sponsor). All patients provided written informed consent.

### Experimental design

Participants saw three different images of electrical devices (a hair dryer, a ventilator, and a toaster) during acquisition, extinction, and test. During acquisition, two out of these three images were associated with an aversive unconditioned stimulus (US), and labeled as “CS+”, while the remaining image was not associated with the US and labeled “CS-” (Figure 1A, top). The association of the images with the US was counterbalanced across subjects. Each image was presented 24 times during acquisition and extinction in a pseudo-randomized fashion, in order to avoid patients anticipating the images and the US (total number of trials in a block: 72). Specifically, a maximum of 3 consecutive presentations of the same item was allowed, irrespective of their particular contingencies. Every cue was presented in one of many different contexts (i.e., videos), which were not related to the type of trial and the occurrence of the US. Four unique context videos were presented during acquisition and another four were presented during extinction. Critically, during extinction, the CS type of one of the items was modified, while the other two remained the same. This led to the distinction between three different types of items: CS++, CS+-, and CS--. After the extinction period, participants performed a test in which they again saw the three items during 16 trials each (total number of trials during the test = 48), which were associated with eight new experimental contexts (the number of contexts was doubled to match those presented during acquisition and extinction). In the test phase, no US was presented for any of the items. The association of the contexts and the cues was completely randomized in every subject, so that contexts were not predictive of the CS. Please note that this approach differs from conventional fear and extinction learning paradigms. Our design was developed to leverage the power of RSA and iEEG to dynamically track the representation of contexts across key hubs of the extinction network. On the other hand, our paradigm aligns with conventional ABC paradigms as it contains three different context categories across experimental phases (See Supplementary Note 4). Similar to the cue-CS associations, the context videos were counterbalanced across subjects and experimental phases. Following previous literature, the reinforcement rate of the CS+ items was set to 50% both during acquisition and extinction, in order to avoid fast associative learning and extinction of the cue CS associations (Dunsmoor et al., 2014).

At every trial, in each experimental phase, participants first saw a fixation cross for 0.5s, and then a context video which was presented for 2s. After the video presentation, a cue was overlayed on top of the video for 1.75s, and then the US was presented for 1s. Finally, a question was presented how much participants had expected a US – i.e., how threatening of safe they had perceived the CS – and participants provided their responses on a scale from 1 (threatening) to 4 (safe).

The US consisted of an unpleasant scream, which was delivered via the computer speakers. The sound levels of the scream were at ∼85 decibels, resulting in a highly aversive but not painful amplitude. The duration of the scream was 1s. Given the 50% reinforcement rate of the US, the scream was presented a total of 36 times in the experiment (CS++: 24 times across acquisition and extinction; CS+- 12 times only during acquisition).

The experiment was programmed in Presentation (Neurobehavioral systems, California, USA), and was deployed on a laptop computer running Microsoft Windows. Patients performed the experiment while sitting in their hospital beds and responded to the memory test using the keyboard of the laptop computer.

Please note that our experimental design is grounded in theoretical considerations from the human fear learning literature and has been adapted to meet the specific requirements of conducting research in implanted epilepsy patients. While its specific combination of features is novel and has not been fully validated, we provide a detailed rationale for each of our design choices, highlighting their similarities and differences to previous research in the field, in Supplementary Note 4.

### Behavioral analysis

We conducted two main behavioral analyses. In the first analysis, we examined average ratings across trials within each experimental phase in order to identify differences in the ratings between phases. A two-way repeated measures ANOVA was used with trial type and experimental phase as factors. Post-hoc tests with Bonferroni correction were applied across conditions after confirming that both a main effect and an interaction were observed (Figure 1). Due to a technical issue during the test phase, one participant was excluded from this analysis, resulting in a sample size of 48 subjects.

In the second analysis, we sorted trials according to their type (CS++, CS+-, and CS--) and compared the ratings for each trial type at every position. Since the comparisons were performed on single-trial data, group level distributions were not normal. Thus, we employed the non-parametric Wilcoxon-signed rank test to assess whether conditions differed at every trial. This was done in order to evaluate the progressive learning of the trial contingencies during acquisition and extinction. In this analysis, we sorted trials by type (CS++, CS+-, and CS--) and performed a Wilcoxon signed-rank test across subjects at each trial to compare performance across conditions. While sorting trials in this way disrupted the absolute trial positions in the experiment, the sequential order of trials within each condition was maintained. For a specific trial (e.g., trial 9), the CS type could be different across subjects. However, after sorting by condition, the first CS++ in all subjects is always compared with the first CS+- and the first CS-- trial, the second with the second, and so on. Thus, the paired Wilcoxon tests were performed across subjects in individual trials corresponding to the same ordinal position in each condition. We corrected for multiple comparisons using cluster-based permutation statistics, shuffling the condition labels at every trial.

### Intracranial EEG recordings

In Paris, recordings were conducted using the ATLAS amplifier (Neuralynx, Bozeman, MO) with a total of 160 channels and a sampling rate of 4096 Hz (Lehongre et al., 2022). In Guangzhou, data was collected using a 256-channel Nihon Kohden Neurofax 1200A Digital System, sampled at 2000 Hz. The number of electrodes and their implantation sites were determined exclusively by clinical requirements. For all analyses, data was downsampled to 1000 Hz. Bipolar referencing was applied, where the activity at each contact point was subtracted from that of the nearest contact on the same electrode, resulting in a total of N-1 virtual channels for an electrode with N channels after re-referencing.

### Channel Localization

Stereotactically implanted electrodes varied in type, number of contacts, and inter-contact distances. In Paris, two types of electrodes were used: standard macro electrodes and macro-micro electrodes. The standard macro electrodes were depth electrodes (Ad-Tech Medical Instrument Corporation, Wisconsin, USA), consisting of 4 to 12 platinum contact points, each with a diameter of 1.12 mm and a length of 2.41 mm, connected via nickel-chromium wiring. The center-to-center distance between adjacent contacts was 5 mm. The macro-micro electrodes (also from AdTech, Wisconsin, USA) were of the Behnke-Fried type, consisting of 8 platinum contact points, each 1.28 mm in diameter and 1.57 mm in length, mounted on the surface of a polyurethane tube with a hollow lumen. In addition, eight 40 µm platinum-iridium microwires were embedded within the macroelectrode, extending 3- 6 mm beyond the tip into the surrounding cerebral tissue (data from these microwires were not analyzed in this study). The inter-contact distance between contacts 1 and 2 was 3 mm, while the distance between subsequent contacts (e.g., 2 and 3, 3 and 4, etc.) was 7 mm.

In Guangzhou, depth electrodes were obtained from Sinovation Medical Technology (Beijing), and consisted of platinum depth electrodes with 7 to 19 contacts each (0.8 mm diameter, 2 mm length, and 1.5 mm spacing).

Electrode localization was conducted separately by our clinical teams in Paris and Guangzhou using very similar methodological approaches. At both sites, post-implantation computed tomography (CT) images were coregistered onto preimplantation magnetic resonance imaging (MRI) data. Both sites used the same software including Freesurfer (http://surfer.nmr.mgh.harvard.edu/), 3DSlicer (https://www.slicer.org), and Statistical Parametric Mapping (SPM; https://www.fil.ion.ucl.ac.uk/spm/).

In Paris, electrode coordinates were extracted using the EPILOC toolbox developed on the STIM platform (Stereotaxy, Techniques, Images, Models; http://pf-stim.cricm.upmc.fr). EPILOC automates several image processing steps using Freesurfer, 3DSlicer, and SPM, complemented by Brainvisa (Rivière et al., 2011). Anatomical models were generated from preoperative MRI data using Freesurfer. Post-operative CT data within the patient’s native space were coregistered onto the preoperative MRI data and then normalized to a Montreal Neurological Institute (MNI) template using SPM. Depth sEEG electrodes were automatically localized in postoperative CT images by segmenting electrode artifacts and classifying them based on their proximity to trajectories planned by stereotactic guidance devices.

In Guangzhou, channel locations were again identified by coregistering the post-implantation CT images to pre-implantation MRIs acquired for each patient. Patient specific anatomical surfaces were generated with Freesurfer and normalized to MNI space using SPM. Electrode locations were determined using 3DSlicer.

A table containing the electrode coordinates in MNI space for all contacts included in the study is presented in Supplementary Table 2. We note that since the implantation scheme is solely determined by clinical criteria, density of sampling within and across brain regions is inherently variable in iEEG research. Several factors might affect the volume of neural tissue sampled, e.g., different electrode configurations, individual implantation schemes, and the selection of ROIs. We performed several control analyses to corroborate that the volume of neural tissue sampled did not differ between our two patient cohorts (see Supplementary Note 5 and Supplementary Figure 2).

### ROI selection

We recorded activity from several brain regions pertaining to the fear learning and extinction network, including the amygdala, hippocampus, the lateral prefrontal cortex (lPFC), the orbitofrontal cortex (OFC) as well as in various areas relevant for processing of sensory stimulus features in the temporal cortex (TMP; Figure 1B). Electrodes from both left and right hemispheres were included in each of the ROIs. We employed the Desikan Killian atlas (Desikan et al., 2006) for label extraction.

Electrodes located at the following locations were labeled as TMP electrodes: ‘inferiortemporal’, ‘middletemporal’, ‘superiortemporal’, ‘transversetemporal’, ‘fusiform’, ‘temporalpole’, ‘bankssts’, ‘parahippocampal’, ‘entorhinal’. This resulted in a total number of 325 electrodes (46 subjects) in TMP.

Electrodes located at the following freesurfer locations were labeled as lPFC electrodes: ‘caudalmiddlefrontal’, ‘parsopercularis’, ‘parsorbitalis’, ‘superiorfrontal’, ‘parstriangularis’, ‘rostralmiddlefrontal’, ‘frontalpole’. This resulted in a total number of 74 electrodes (23 subjects) in lPFC.

Since the ROIs lPFC and TMP cover relatively large brain areas, we provide information about electrode localizations in their specific subregions in Supplementary Figure 3.

Electrodes located at the following freesurfer locations were labeled as OFC electrodes: ‘lateralorbitofrontal’, ‘medialorbitofrontal’. This resulted in a total number of 35 electrodes (12 subjects) in OFC.

In the AMY, we recorded from 82 electrodes in 32 subjects. In the HPC, we recorded from 115 electrodes in 34 subjects. A video depiction of AMY and HPC electrode locations is provided in Supplementary video 1.

In all patients, we removed channels located in white matter, resulting in 631 clean channels across all patients (12.62 ± 6.66 channels per patient).

### Preprocessing

Preprocessing was performed on the entire raw data using EEGLAB (Delorme & Makeig, 2004), and included high-pass filtering at a frequency of 0.1Hz and low-pass filtering at a frequency of 200Hz. We also applied a band-stop (notch) filter with frequencies of 49-51Hz, 99-101Hz, and 149-151Hz. Artifact rejection was performed using a previously published method (Staresina et al., 2015). This automatic detection algorithm identifies epileptiform activity based on three key criteria: (1) the amplitude of the time-series, (2) the gradient (i.e., the amplitude difference between adjacent time points), and (3) the amplitude of the data after applying a 250-Hz high-pass filter, which further enhances the detection of epileptogenic spikes. Each time point in the raw iEEG data was converted into a z-score based on the participant- specific mean and standard deviation of these three measures across the whole experiment in each channel independently. A time point was marked as artifactual if it exceeded a z-score of 6 in any measure or met a conjunction threshold of an amplitude z-score of 4 combined with either a gradient or high-frequency z-score of 3.

We found that the combination of raw EEG amplitudes, EEG gradients, and high-pass filtered EEG amplitudes at 250 Hz provided high sensitivity for detecting epileptiform activity and other artifacts. This was further validated through manual identification of artifacts in randomly selected subsets of the data. To ensure that all epileptic activity was completely removed from our data, we not only excluded the detected time points but also removed the 1,000ms preceding and following each detected artifact sample.

After identifying and marking the segments of the data contaminated with artifacts, we segmented the data into 7-second epochs (from -3 to 4 s) around the presentation of each CS cue. Any epoch containing artifact segments detected in the continuous (non-epoched) data was entirely removed from further analysis. Additionally, we visually examined raw time-series plots and spectrograms to assess the presence of artifacts in the frequency domain. The number of epochs removed due to artifacts varied depending on the signal quality of each subject and was at 20.6 ± 19.2 epochs of cue presentation across subjects and channels. The number of excluded trials in each condition of the experiment is presented in the Supplementary Table 3.

### Time-frequency analysis

Using the FieldTrip toolbox (Oostenveld et al., 2011), we decomposed the signal using complex Morlet wavelets with a variable number of cycles, i.e., linearly increasing in 29 steps between 3 cycles (at 1 Hz) and 6 cycles (at 29 Hz) for the low-frequency range, and in 15 steps from 6 cycles (at 30 Hz) to 12 cycles (at 100LJHz) for the high-frequency range (Pacheco Estefan et al., 2019; Staresina et al., 2016). The resulting time-series of frequency-specific power values were then z-scored by taking as a reference the mean activity across all trials in the experiment (Fellner et al., 2020). This type of normalization was applied to remove any common feature of the signal unrelated to the stability of the cue representations or the encoding of context-specific information.

We contrasted oscillatory power values in the frequency ranges of interest between CS+ and CS- trials during acquisition and extinction (Figure 2A). Note that in the analysis during extinction, the data for the CS- condition during extinction includes both CS+- and CS-- trials (i.e., CS- trials that were either CS+ or CS- during acquisition). This was done to focus on the current valence of the CS during extinction—which we assumed would be the primary factor influencing theta power— and to maximize statistical power by including all available trials in the CS- condition. The resulting time-frequency data (with the same decomposition parameters) was employed in our pattern similarity analyses (see below).

We only included subjects with at least 8 trials in the group comparisons, resulting in a total of N = 32 subjects in the analysis performed in the amygdala. The number of CS+ trials was 38.72 ± 8.48 (Mean ± STD) during acquisition and 19.46 ± 4.57 during extinction in this region.

### Contrast-based representational similarity analysis

We employed temporally resolved Representational Similarity Analysis (RSA) to evaluate the dynamics of context and cue representations following previous work (Pacheco Estefan et al., 2019, 2021). We assessed the stability of neural representations by computing the within-item similarity of each cue (i.e., the average similarity for all repeated presentations of the same item, Figure 3A). We compared this metric between CS+ and CS- trials, i.e., tested whether the repeated visual presentation of a cue associated with a US (CS+ items) elicited more stable response patterns as compared to the repeated presentation of non-predictive cues (CS- items; Figure 3A). This was done separately during acquisition and extinction. The CS+ vs CS- comparisons were performed using paired t-tests after Fisher Z-transforming the rho values obtained in each trial and condition, in each subject independently. Note that the computation of item-stability implies that representational patterns across repeated presentations of the same items cannot be averaged as is often done in RSA.

In addition, we investigated the presence of context-specific representations by comparing the similarity of same-contexts versus different-contexts in each of our ROIs. We assessed similarities for same-context and different-context item pairs and averaged across all combinations of items in the same condition in each subject independently. The same-context and different-context correlations were then statistically compared at the group level using *t*-tests (rho values were Fisher z-transformed before contrasting them). This metric was also computed during acquisition and extinction. Similar to the item stability metric previously described, we did not average the representational patterns corresponding to the repeated presentation of the same context video but performed the similarity comparisons across all instances in which a particular video was presented. In some of our context-specific analysis (presented in Figure 3), we locked the data to the presentation of the cue to facilitate the comparison with item stability results. Note that both cues and videos are presented simultaneously during the cue presentation (the cue is overlayed on top of the video), but the videos are additionally presented separately for two seconds before the cues.

Representational patterns were defined by specifying a 500ms time window, in which averaged time courses of frequency-specific power values (all frequencies in the 1-100Hz range) were included across all contacts in the respective ROI. A representational pattern was thus composed of activity of N electrodes x 44 frequencies in each 500ms window. Note that the number of channels included varied depending on the number of electrodes available for a particular subject/ROI. These two-dimensional arrays were concatenated into 1D vectors for similarity comparisons. In total, the number of electrodes included in each of our ROIs was the following (Mean ± STD across subjects): TMP: 7.06 ± 4.23; Amygdala: 2.56 ± 1.39; Hippocampus: 3.38 ± 1.82; OFC: 2.91 ± 0.9; lPFC: 3.21 ± 1.83.

Pairwise (Spearman) correlations among representational patterns were computed in sliding time windows (windows of 500ms computed at every 50ms, i.e., with a 90% overlap), resulting in a time series of neural representational similarity matrices (RSMs) for each of our ROIs. In the main analyses (Figure 3 and 4), we assessed the correlation of these representational patterns across all matching time-points, resulting in a time-series of item stability or context specificity values for each condition. The time-series were averaged in each condition for each subject independently (after Fisher z-transforming them), and the resulting average time-series were contrasted via paired *t*-tests across conditions at the group level. Multiple comparisons correction was performed using cluster-based permutation statistics (see below)

The number of trials for CS+ and CS- items was unbalanced during acquisition and extinction (two out of three items were labeled as CS+ during acquisition and only one as CS-, while the reverse was true during extinction). We thus computed within-item similarity separately for each of the two items with the same contingency in a given phase, and then averaged the results before contrasting the time series with the time-series of the item corresponding to the opposite CS.

We only included subjects with at least 8 trials in each condition, leading to the following number of participants for the item stability analysis: AMY: 32; HPC: 34; OFC: 12; lPFC: 22 for both acquisition and extinction. In the TMP, the number of subjects for the item stability analysis was 43 during acquisition and 44 during extinction. The number of participants included in the context specificity analysis was the same in both experimental phases in all regions: lPFC: 22; AMY: 32; HPC: 34; OFC: 12; TMP: 46.

In all pattern similarity plots, correlations corresponding to each 500ms window were assigned to the time point at the center of the respective window (e.g., a time bin corresponding to activity from 0 to 500ms was assigned to 250ms).

### Coordination analyses

We computed a single-trial metric of item-stability by assessing the similarity of the neural representation of an item presented in a trial with the representation of the same item in all other trials during a particular experimental phase. In addition, we computed a single-trial metric of context specificity by subtracting the similarity of each context video with all repeated presentations of the same video and the similarity with all different videos in a particular experimental phase. These metrics were computed in each of our ROIs and then correlated between different ROIs across trials for each participant. Correlations were calculated in sliding time windows (500ms windows, overlapping by 95%) across all possible combinations of time points. This resulted in one correlation value for each subject in each time-bin x time-bin pair. We assessed potential temporal offsets because the latency and duration of representations may differ across brain regions, making it more likely that coordination is temporally shifted rather than simultaneous (see King & Dehaene, 2014; Pacheco Estefan et al., 2019). Notably, the relative timing of coordination offers insights into the temporal order of representations across brain regions during fear extinction. Statistical significance was assessed by contrasting these rho values against zero at the group level, and correction for multiple comparisons was performed using cluster-based permutation statistics (see below).

To confirm that the trial-level results were not driven by the main condition differences in the representational feature vectors representing CS+ and CS- items, we performed all trial-level correlation analyzes after separately Z-scoring the reinstatement values in CS+ and CS- trials. We obtained equivalent results in all of reported effects. In additional analyses, we assessed connectivity between ROIs where we observed significant coordination effects using the weighted phase lag index (wPLI). Results are presented in Supplementary Note 6.

The number of subjects included in the trial-based RSA analyses depended on two criteria: 1) the number of trials in each condition, which was set to a minimum of 8 trials, consistent with the contrast-based RSA analyses (see above). A subject was excluded from this analysis if the number of trials was below 8 in any of the conditions tested, and 2) the number of subjects with electrodes in both ROIs, which was based solely on clinical needs. The number of participants included in the trial-based analyses comparing item stability between regions was the following: TMP- AMY: 28; TMP-HPC: 31; TMP-lPFC: 18; TMP-OFC; 11; AMY-HPC: 24; AMY-lPFC: 14; AMY-OFC: 8. The number of participants included in the trial-based comparisons of lPFC context specificity with context specificity in the other areas was the following: lPFC-AMY: 14; lPFC-TMP: 18; lPFC-HPC: 15; lPFC-OFC: 11.

### Reinstatement analysis

To investigate whether lPFC context specificity during extinction predicted reinstatement in the TMP during the test phase, we conducted a subject-level correlation analysis. First, we calculated a metric of context specificity in the lPFC by averaging the difference between same-context and different-context correlations within the time period where significant context effects were observed in this region during extinction (from 0.8s-1.15s; Figure 4C, middle). These correlations were averaged both across trials and over time within this time window, resulting in a single value of lPFC context specificity for each subject. Second, we calculated reinstatement across experimental phases, comparing activity patterns during the test with those observed either during acquisition or during extinction (i.e., acquisition-test reinstatement and extinction-test reinstatement). We specifically focused on the TMP, where the main condition differences were observed (i.e., higher item stability for CS+ as compared to CS- trials). We correlated each trial during acquisition and extinction with each trial during test in each subject independently. We then averaged these correlations across trials and across time (focusing on the same time window of significant lPFC context specificity) and subtracted acquisition-test reinstatement and extinction-test reinstatement, resulting in a single value of TMP reinstatement for each subject. Finally, we correlated context specificity and reinstatement across subjects using Spearman correlations (Figure 6B).

To further investigate whether the observed correlation between the lPFC and TMP was unique to the period of context specificity in the lPFC, we conducted a temporally resolved analysis. Employing sliding time windows of 500ms with a 95% overlap, we again correlated same and different items during both acquisition and extinction and quantified the time course of item-specific reinstatement during the test phase. In the lPFC, we selected context-specific representations in the same cluster used in the primary analysis (described above), and correlated these values across subjects at every time point of the TMP reinstatement time course using Spearman correlations. This approach yielded a time course of Rho and p-values, which we subsequently corrected for multiple comparisons using cluster-based permutation statistics (see below).

### Functional relevance of extinction reinstatement

To assess the functional significance of reinstatement during the test phase, we computed a trial-level metric of reinstatement values. Specifically, we measured the similarity of each trial in the test phase to the same and different items from the extinction phase and subtracted these two values. This was conducted at matching time points within each trial, generating a time course of item reinstatement for every trial. We focused on the TMP, and specifically on the time period when significant differences in item stability between CS+ and CS- trials were detected. We averaged reinstatement values in this window, resulting in a trial-level metric of item-specific reinstatement for each subject. These reinstatement values were then correlated with the participants’ subjective ratings using Spearman correlations (after Fisher Z-transforming them). The resulting values were tested against zero at the group level to determine statistical significance (Figure 7).

In our experiment, the average number of trials in which participants failed to provide a response was considerable (12.65 ± 15.47 per subject), primarily because responding was not a prerequisite for progressing to the next trial in any experimental phase. Consequently, we applied a more liberal threshold for the minimum number of trials included in this analysis, setting it at 4 instead of 8 trials. We repeated the analysis with the stricter threshold of 8 trials and found qualitatively similar results (CS++: t(14)= 2.17; p = 0.048; CS+-: t(14)= 2.29; p = 0.037; CS--: t(13)= -0.67; p = 0.51).

To account for potential group-level biases introduced by outlier effects, we repeated our analysis after excluding trials with TMP reinstatement values exceeding 2 standard deviations from the mean for each subject. On average, 7.2 ± 4.9 trials were excluded per subject, resulting in a final sample of 17 subjects. The results remained largely consistent (CS++: t(17) = 0.849, p = 0.408; CS+-: t(16) = 2.51, p = 0.0231; CS--: t(18) = -0.0126, p = 0.99).

### Multiple comparisons corrections

We performed cluster-based permutation statistics to correct for multiple comparisons in the oscillatory power analyses (Figure 2), the contrast-based pattern similarity analyses (Figures 3B, 3D, 4B and 4C), the trial-based pattern similarity analysis (Figures 3F and 4D), and the subject-level correlation analysis between lPFC and TMP (Figure 6).

In the oscillatory power analyses, we contrasted CS+ and CS- time-frequency maps using paired t-tests, after shuffling the trial labels 1,000 times in each subject independently. We considered significant a time-point if the difference between these surrogate conditions was significant at *p* < 0.05 (two-tailed tests were employed). At every permutation, we computed clusters of significant values defined as contiguous regions in time-frequency space where significant correlations were observed. We took the largest cluster at each permutation to build the null distribution. We only considered significant those contiguous time pairs in the empirical (non-shuffled) data whose summed *t*-values exceeded the summed *t*-value of 95% of the surrogate clusters (corresponding to a corrected P < 0.05; see Maris & Oostenveld, 2007).

Since the contrast-based pattern similarity analyses were based on matching time points, we contrasted time-series of different conditions (CS+ vs CS- items or same vs different contexts) at different time-points using *t*-tests, after shuffling the trial labels of each condition 1,000 times in each subject independently. Adjacent regions of significant differences were formed along one dimension only (time), resulting in a distribution of surrogate *t*-values under the assumption of the null hypothesis. Similar to the procedure employed in the oscillatory power analysis, we only considered significant those contiguous time pairs in the empirical data whose summed *t*-values were above the summed *t*-value of 95% of the distribution of surrogate clusters (Maris & Oostenveld, 2007).

In the trial-based pattern similarity analysis, a temporal generalization approach was employed, and therefore clusters of significant regions were formed in 2 dimensions (time x time). Item stability or context specificity was correlated across trials between brain regions at the group level using paired t-tests as in the original data. T-values of adjacent regions where significant differences were observed were summed. At every permutation, we shuffled the trial labels in one of the regions and performed the analysis again, storing the cluster with the highest absolute t-value. This resulted in a distribution of surrogate t-values under the null hypothesis of no condition differences. Similar to the procedure we employed in the contrast-based analyses, we then computed p-values by ranking the observed t-values of the biggest cluster in the empirical data with this surrogate distribution. Time periods with contiguous significant regions exceeding 95% of surrogate clusters were considered significant (Maris & Oostenveld, 2007).

In the temporally-resolved subject-level correlation analysis (Figure 6D), we identified clusters of contiguous subject-level correlations after shuffling the lPFC and TMP subject IDs 1,000 times. This approach yielded a total of 1,000 time courses of Rho and p-values. Rho values were summed in the biggest cluster of contiguous significant correlations at every permutation, resulting in a distribution of Rho values expected under the null hypothesis. We then ranked the summed rho values within this cluster in the empirical data with respect to the surrogate distribution to determine statistical significance.

In addition to cluster-based permutations, we also corrected our results for multiple comparisons using the Bonferroni method. Given that we included 5 ROIs, the alpha level for significance was set to p = 0.05/5 = 0.01.

## Acknowledgements

This work was supported by the Deutsche Forschungsgemeinschaft (DFG, German Research Foundation) through grant SFB 1280 (projects A01 and A02) project number 316803389.

## Data Availability

Anonymized intracranial EEG data supporting the findings of this study will be made available upon publication.

## Code Availability

Custom-written Matlab code supporting the findings of this study will be made available upon publication.

## Contributions

Conceptualization, M-C.F., D.P.-E. and N.A.; methodology, D.P.-E., A.B., G.J., and N.A.; data collection, M.-C.F., K.L., V.L., V.F. and J.Y.; data analysis: D.P.-E. and N.A.; writing—original draft, D.P.-E.; writing—review and editing, D.P.-E. and N.A.; funding acquisition, N.A., O.G.; resources, V.N., O.G., L.S., B.H., Q.C.; supervision N.A.

## Notes

### Competing Interest Statement

The authors have declared no competing interest.

## References

Axmacher, N. (2023). Intracranial EEG: A Guide for Cognitive Neuroscientists. Springer International Publishing.

Balooch, S. B., Neumann, D. L., & Boschen, M. J. (2012). Extinction treatment in multiple contexts attenuates ABC renewal in humans. Behaviour Research and Therapy, 50(10), 604–609. 10.1016/j.brat.2012.06.003

Bandarian Balooch, S., & Neumann, D. L. (2011). Effects of multiple contexts and context similarity on the renewal of extinguished conditioned behaviour in an ABA design with humans. Learning and Motivation, 42(1), 53–63. 10.1016/j.lmot.2010.08.008

Beckers, T., Hermans, D., Lange, I., Luyten, L., Scheveneels, S., & Vervliet, B. (2023). Understanding clinical fear and anxiety through the lens of human fear conditioning. Nature Reviews Psychology, 2(4), 233–245. 10.1038/s44159-023-00156-1

Bierwirth, P., Antov, M. I., & Stockhorst, U. (2023). Oscillatory and non-oscillatory brain activity reflects fear expression in an immediate and delayed fear extinction task. Psychophysiology, 60(8), e14283. 10.1111/psyp.14283

Bierwirth, P., Sperl, M. F. J., Antov, M. I., & Stockhorst, U. (2021). Prefrontal Theta Oscillations Are Modulated by Estradiol Status During Fear Recall and Extinction Recall. Biological Psychiatry: Cognitive Neuroscience and Neuroimaging, 6(11), 1071–1080. 10.1016/j.bpsc.2021.02.011

Bocchio, M., Nabavi, S., & Capogna, M. (2017). Synaptic Plasticity, Engrams, and Network Oscillations in Amygdala Circuits for Storage and Retrieval of Emotional Memories. Neuron, 94(4), 731–743. 10.1016/j.neuron.2017.03.022

Boddez, Y., Baeyens, F., Luyten, L., Vansteenwegen, D., Hermans, D., & Beckers, T. (2013). Rating data are underrated: Validity of US expectancy in human fear conditioning. Journal of Behavior Therapy and Experimental Psychiatry, 44(2), 201–206. 10.1016/j.jbtep.2012.08.003

Bouton, M. E. (2004). Context and behavioral processes in extinction. *Learning & Memory (Cold Spring Harbor*, N.Y.), 11(5), 485–494. 10.1101/lm.78804

Bouton, M. E., Maren, S., & McNally, G. P. (2021). BEHAVIORAL AND NEUROBIOLOGICAL MECHANISMS OF PAVLOVIAN AND INSTRUMENTAL EXTINCTION LEARNING. Physiological Reviews, 101(2), 611–681. 10.1152/physrev.00016.2020

Bouton, M. E., & Swartzentruber, D. (1991). Sources of relapse after extinction in Pavlovian and instrumental learning. Clinical Psychology Review, 11(2), 123–140.

Burke, J. F., Long, N. M., Zaghloul, K. A., Sharan, A. D., Sperling, M. R., & Kahana, M. J. (2014). Human intracranial high-frequency activity maps episodic memory formation in space and time. New Horizons for Neural Oscillations, 85, 834–843. 10.1016/j.neuroimage.2013.06.067

Cattani, A., Arnold, D. B., McCarthy, M., & Kopell, N. (2024). Basolateral amygdala oscillations enable fear learning in a biophysical model. eLife, 12, RP89519. 10.7554/eLife.89519

Chen, S., Tan, Z., Xia, W., Gomes, C. A., Zhang, X., Zhou, W., Liang, S., Axmacher, N., & Wang, L. (2021). Theta oscillations synchronize human medial prefrontal cortex and amygdala during fear learning. Science Advances, 7(34), eabf4198. 10.1126/sciadv.abf4198

Constantinou, E., Purves, K. L., McGregor, T., Lester, K. J., Barry, T. J., Treanor, M., Craske, M. G., & Eley, T. C. (2021). Measuring fear: Association among different measures of fear learning. Journal of Behavior Therapy and Experimental Psychiatry, 70, 101618. 10.1016/j.jbtep.2020.101618

Corcoran, K. A., & Maren, S. (2004). Factors regulating the effects of hippocampal inactivation on renewal of conditional fear after extinction. Learning & Memory, 11(5), 598–603.

Corneille, O., & Gawronski, B. (2024). Self-reports are better measurement instruments than implicit measures. Nature Reviews Psychology, 3(12), 835–846. 10.1038/s44159-024-00376-z

Costa, M., Lozano-Soldevilla, D., Gil-Nagel, A., Toledano, R., Oehrn, C. R., Kunz, L., Yebra, M., Mendez-Bertolo, C., Stieglitz, L., Sarnthein, J., Axmacher, N., Moratti, S., & Strange, B. A. (2022). Aversive memory formation in humans involves an amygdala-hippocampus phase code. Nature Communications, 13(1), 6403. 10.1038/s41467-022-33828-2

Costa, M., Pacheco, D., Gil-Nagel, A., Toledano, R., Imbach, L., Sarnthein, J., & Strange, B. A. (2024). Retrieval of human aversive memories involves reactivation of gamma activity patterns in the hippocampus that originate in the amygdala during encoding. bioRxiv, 2024.01.18.576178. 10.1101/2024.01.18.576178

Courtin, J., Chaudun, F., Rozeske, R. R., Karalis, N., Gonzalez-Campo, C., Wurtz, H., Abdi, A., Baufreton, J., Bienvenu, T. C. M., & Herry, C. (2014). Prefrontal parvalbumin interneurons shape neuronal activity to drive fear expression. Nature, 505(7481), 92–96. 10.1038/nature12755

Craske, M. G., Treanor, M., Conway, C. C., Zbozinek, T., & Vervliet, B. (2014). Maximizing exposure therapy: An inhibitory learning approach. Behaviour Research and Therapy, 58, 10–23. 10.1016/j.brat.2014.04.006

Darna, M., Stolz, C., Jauch, H.-S., Strumpf, H., Hopf, J.-M., Seidenbecher, C. I., Schott, B. H., & Richter, A. (2024). Frontal Theta Oscillations and Cognitive Flexibility: Age-Related Modulations in EEG Activity. bioRxiv, 2024.07.05.602082. 10.1101/2024.07.05.602082

Delorme, A., & Makeig, S. (2004). EEGLAB: an open source toolbox for analysis of single-trial EEG dynamics including independent component analysis. Journal of Neuroscience Methods, 134(1), 9–21. 10.1016/j.jneumeth.2003.10.009

Desikan, R. S., Ségonne, F., Fischl, B., Quinn, B. T., Dickerson, B. C., Blacker, D., Buckner, R. L., Dale, A. M., Maguire, R. P., Hyman, B. T., Albert, M. S., & Killiany, R. J. (2006). An automated labeling system for subdividing the human cerebral cortex on MRI scans into gyral based regions of interest. NeuroImage, 31(3), 968–980. 10.1016/j.neuroimage.2006.01.021

Dunsmoor, J. E., Ahs, F., Zielinski, D. J., & LaBar, K. S. (2014). Extinction in multiple virtual reality contexts diminishes fear reinstatement in humans. Extinction, 113, 157–164. 10.1016/j.nlm.2014.02.010

Eichenbaum, H. (2017a). Memory: Organization and Control. Annual Review of Psychology, 68, 19–45. 10.1146/annurev-psych-010416-044131

Eichenbaum, H. (2017b). Prefrontal–hippocampal interactions in episodic memory. Nature Reviews Neuroscience, 18(9), 547–558. 10.1038/nrn.2017.74

Engen, H. G., & Anderson, M. C. (2018). Memory Control: A Fundamental Mechanism of Emotion Regulation. Trends in Cognitive Sciences, 22(11), 982–995. 10.1016/j.tics.2018.07.015

Fell, J., & Axmacher, N. (2011). The role of phase synchronization in memory processes. Nature Reviews. Neuroscience, 12(2), 105–118. 10.1038/nrn2979

Fell, J., Ludowig, E., Staresina, B. P., Wagner, T., Kranz, T., Elger, C. E., & Axmacher, N. (2011). Medial Temporal Theta/Alpha Power Enhancement Precedes Successful Memory Encoding: Evidence Based on Intracranial EEG. The Journal of Neuroscience, 31(14), 5392. 10.1523/JNEUROSCI.3668-10.2011

Fellner, M. C., Waldhauser, G. T., & Axmacher, N. (2020). Tracking Selective Rehearsal and Active Inhibition of Memory Traces in Directed Forgetting. Current Biologyl: CB, 30(13), 2638–2644.e4. 10.1016/j.cub.2020.04.091

Fenton, G. E., Halliday, D. M., Mason, R., & Stevenson, C. W. (2014). Medial prefrontal cortex circuit function during retrieval and extinction of associative learning under anesthesia. Neuroscience, 265, 204–216. 10.1016/j.neuroscience.2014.01.028

Foster, B. L., Dastjerdi, M., & Parvizi, J. (2012). Neural populations in human posteromedial cortex display opposing responses during memory and numerical processing. Proceedings of the National Academy of Sciences, 109(38), 15514–15519. 10.1073/pnas.1206580109

Foster, B. L., Rangarajan, V., Shirer, W. R., & Parvizi, J. (2015). Intrinsic and Task-Dependent Coupling of Neuronal Population Activity in Human Parietal Cortex. Neuron, 86(2), 578–590. 10.1016/j.neuron.2015.03.018

Fullana, M. A., Albajes-Eizagirre, A., Soriano-Mas, C., Vervliet, B., Cardoner, N., Benet, O., Radua, J., & Harrison, B. J. (2018). Fear extinction in the human brain: A meta-analysis of fMRI studies in healthy participants. Neuroscience & Biobehavioral Reviews, 88, 16–25. 10.1016/j.neubiorev.2018.03.002

Fullana, M. A., Harrison, B. J., Soriano-Mas, C., Vervliet, B., Cardoner, N., Àvila-Parcet, A., & Radua, J. (2016). Neural signatures of human fear conditioning: An updated and extended meta-analysis of fMRI studies. Molecular Psychiatry, 21(4), 500–508. 10.1038/mp.2015.88

Garfinkel, S. N., Abelson, J. L., King, A. P., Sripada, R. K., Wang, X., Gaines, L. M., & Liberzon, I. (2014). Impaired Contextual Modulation of Memories in PTSD: An fMRI and Psychophysiological Study of Extinction Retention and Fear Renewal. The Journal of Neuroscience, 34(40), 13435. 10.1523/JNEUROSCI.4287-13.2014

Gattas, S., Larson, M. S., Mnatsakanyan, L., Sen-Gupta, I., Vadera, S., Swindlehurst, A. L., Rapp, P. E., Lin, J. J., & Yassa, M. A. (2023). Theta mediated dynamics of human hippocampal-neocortical learning systems in memory formation and retrieval. Nature Communications, 14(1), 8505. 10.1038/s41467-023-44011-6

Gilmartin, M. R., Balderston, N. L., & Helmstetter, F. J. (2014). Prefrontal cortical regulation of fear learning. Trends in Neurosciences, 37(8), 455–464. 10.1016/j.tins.2014.05.004

Griffiths, B. J., Parish, G., Roux, F., Michelmann, S., van der Plas, M., Kolibius, L. D., Chelvarajah, R., Rollings, D. T., Sawlani, V., Hamer, H., Gollwitzer, S., Kreiselmeyer, G., Staresina, B., Wimber, M., & Hanslmayr, S. (2019). Directional coupling of slow and fast hippocampal gamma with neocortical alpha/beta oscillations in human episodic memory. Proceedings of the National Academy of Sciences, 116(43), 21834–21842. 10.1073/pnas.1914180116

Hennings, A. C., McClay, M., Drew, M. R., Lewis-Peacock, J. A., & Dunsmoor, J. E. (2022). Neural reinstatement reveals divided organization of fear and extinction memories in the human brain. Current Biology, 32(2), 304–314.e5. 10.1016/j.cub.2021.11.004

Herweg, N. A., Solomon, E. A., & Kahana, M. J. (2020). Theta Oscillations in Human Memory. Trends in Cognitive Sciences, 24(3), 208–227. 10.1016/j.tics.2019.12.006

Holland, P. C., & Bouton, M. E. (1999). Hippocampus and context in classical conditioning. Current Opinion in Neurobiology, 9(2), 195–202.

Jacobs, J. (2014). Hippocampal theta oscillations are slower in humans than in rodents: Implications for models of spatial navigation and memory. Philosophical Transactions of the Royal Society B: Biological Sciences, 369(1635), 20130304. 10.1098/rstb.2013.0304

Karalis, N., Dejean, C., Chaudun, F., Khoder, S., Rozeske, R. R., Wurtz, H., Bagur, S., Benchenane, K., Sirota, A., Courtin, J., & Herry, C. (2016). 4-Hz oscillations synchronize prefrontal–amygdala circuits during fear behavior. Nature Neuroscience, 19(4), 605–612. 10.1038/nn.4251

King, J.-R., & Dehaene, S. (2014). Characterizing the dynamics of mental representations: The temporal generalization method. Trends in Cognitive Sciences, 18(4), 203–210.

Kinner, V. L., Merz, C. J., Lissek, S., & Wolf, O. T. (2016). Cortisol disrupts the neural correlates of extinction recall. NeuroImage, 133, 233–243. 10.1016/j.neuroimage.2016.03.005

Knight, D. C., Nguyen, H. T., & Bandettini, P. A. (2003). Expression of conditional fear with and without awareness. Proceedings of the National Academy of Sciences, 100(25), 15280–15283. 10.1073/pnas.2535780100

Knight, D. C., Waters, N. S., & Bandettini, P. A. (2009). Neural substrates of explicit and implicit fear memory. NeuroImage, 45(1), 208–214. 10.1016/j.neuroimage.2008.11.015

Kobelt, M., Waldhauser, G., Rupietta, A., Heinen, R., Rau, E., Kessler, H., & Axmacher, N. (2024). The memory trace of an intrusive trauma-analog episode. Current Biology, 34(8), 1657–1669.

Kriegeskorte, N., & Diedrichsen, J. (2019). Peeling the Onion of Brain Representations. Annual Review of Neuroscience, 42, 407–432. 10.1146/annurev-neuro-080317-061906

Kriegeskorte, N., Mur, M., & Bandettini, P. A. (2008). Representational similarity analysis-connecting the branches of systems neuroscience. Frontiers in Systems Neuroscience, 4.

Lebois, L. A., Seligowski, A. V., Wolff, J. D., Hill, S. B., & Ressler, K. J. (2019). Augmentation of extinction and inhibitory learning in anxiety and trauma-related disorders. Annual Review of Clinical Psychology, 15(1), 257–284.

Lega, B. C., Jacobs, J., & Kahana, M. (2012). Human hippocampal theta oscillations and the formation of episodic memories. Hippocampus, 22(4), 748–761. 10.1002/hipo.20937

Lehongre, K., Lambrecq, V., Whitmarsh, S., Frazzini, V., Cousyn, L., Soleil, D., Fernandez-Vidal, S., Mathon, B., Houot, M., Lemaréchal, J.-D., Clemenceau, S., Hasboun, D., Adam, C., & Navarro, V. (2022). Long-term deep intracerebral microelectrode recordings in patients with drug-resistant epilepsy: Proposed guidelines based on 10-year experience. NeuroImage, 254, 119116. 10.1016/j.neuroimage.2022.119116

Lesting, J., Daldrup, T., Narayanan, V., Himpe, C., Seidenbecher, T., & Pape, H.-C. (2013). Directional Theta Coherence in Prefrontal Cortical to Amygdalo-Hippocampal Pathways Signals Fear Extinction. PLOS ONE, 8(10), e77707. 10.1371/journal.pone.0077707

Lesting, J., Narayanan, R. T., Kluge, C., Sangha, S., Seidenbecher, T., & Pape, H.-C. (2011). Patterns of Coupled Theta Activity in Amygdala-Hippocampal-Prefrontal Cortical Circuits during Fear Extinction. PLOS ONE, 6(6), e21714. 10.1371/journal.pone.0021714

Likhtik, E., Stujenske, J. M., A Topiwala, M., Harris, A. Z., & Gordon, J. A. (2014). Prefrontal entrainment of amygdala activity signals safety in learned fear and innate anxiety. Nature Neuroscience, 17(1), 106–113. 10.1038/nn.3582

Lissek, S., Biggs, A. L., Rabin, S. J., Cornwell, B. R., Alvarez, R. P., Pine, D. S., & Grillon, C. (2008). Generalization of conditioned fear-potentiated startle in humans: Experimental validation and clinical relevance. Behaviour Research and Therapy, 46(5), 678–687. 10.1016/j.brat.2008.02.005

Lissek, S., & Tegenthoff, M. (2024). Dissimilarities of neural representations of extinction trials are associated with extinction learning performance and renewal level. Frontiers in Behavioral Neuroscience, 18, 1307825.

Liu, J., Zhang, H., Yu, T., Ren, L., Ni, D., Yang, Q., Lu, B., Zhang, L., Axmacher, N., & Xue, G. (2021). Transformative neural representations support long-term episodic memory. Science Advances, 7(41), eabg9715.

Liu, X., Ramirez, S., Pang, P. T., Puryear, C. B., Govindarajan, A., Deisseroth, K., & Tonegawa, S. (2012). Optogenetic stimulation of a hippocampal engram activates fear memory recall. Nature, 484(7394), 381–385. 10.1038/nature11028

Liu, Y., Ye, S., Li, X.-N., & Li, W.-G. (2024). Memory Trace for Fear Extinction: Fragile yet Reinforceable. Neuroscience Bulletin, 40(6), 777–794. 10.1007/s12264-023-01129-3

Lonsdorf, T. B., Menz, M. M., Andreatta, M., Fullana, M. A., Golkar, A., Haaker, J., Heitland, I., Hermann, A., Kuhn, M., Kruse, O., Meir Drexler, S., Meulders, A., Nees, F., Pittig, A., Richter, J., Römer, S., Shiban, Y., Schmitz, A., Straube, B., … Merz, C. J. (2017). Don’t fear ‘fear conditioning’: Methodological considerations for the design and analysis of studies on human fear acquisition, extinction, and return of fear. Neuroscience & Biobehavioral Reviews, 77, 247–285. 10.1016/j.neubiorev.2017.02.026

Lu, Y., Wang, C., Chen, C., & Xue, G. (2015). Spatiotemporal Neural Pattern Similarity Supports Episodic Memory. Current Biology, 25(6), 780–785. 10.1016/j.cub.2015.01.055

Ma, Y., Kyuchukova, D., Jiao, F., Batsikadze, G., Nitsche, M. A., & Yavari, F. (2025). The impact of temporal distribution on fear extinction learning. International Journal of Clinical and Health Psychology, 25(1), 100536.

Maguire, E. A., & Mullally, S. L. (2013). The hippocampus: A manifesto for change. Journal of Experimental Psychology: General, 142(4), 1180.

Manning, J. R. (2023). How Can I Identify Stimulus-Driven Neural Activity Patterns in Multi-Patient ECoG Data? In N. Axmacher (Ed.), Intracranial EEG: A Guide for Cognitive Neuroscientists (pp. 803–836). Springer International Publishing. 10.1007/978-3-031-20910-9_48

Maren, S., Phan, K. L., & Liberzon, I. (2013). The contextual brain: Implications for fear conditioning, extinction and psychopathology. Nature Reviews Neuroscience, 14(6), 417–428. 10.1038/nrn3492

Maren, S., & Quirk, G. J. (2004). Neuronal signalling of fear memory. Nature Reviews Neuroscience, 5(11), 844–852. 10.1038/nrn1535

Maris, E., & Oostenveld, R. (2007). Nonparametric statistical testing of EEG- and MEG-data. Journal of Neuroscience Methods, 164(1), 177–190. 10.1016/j.jneumeth.2007.03.024

McClelland, J. L., McNaughton, B. L., & O’Reilly, R. C. (1995). Why there are complementary learning systems in the hippocampus and neocortex: Insights from the successes and failures of connectionist models of learning and memory. Psychological Review, 102(3), 419–457. 10.1037/0033-295X.102.3.419

McNally, R. J. (2007). Mechanisms of exposure therapy: How neuroscience can improve psychological treatments for anxiety disorders. New Approaches to the Study of Change in Cognitive Behavioral Therapies, 27(6), 750–759. 10.1016/j.cpr.2007.01.003

Méndez-Bértolo, C., Moratti, S., Toledano, R., Lopez-Sosa, F., Martínez-Alvarez, R., Mah, Y. H., Vuilleumier, P., Gil-Nagel, A., & Strange, B. A. (2016). A fast pathway for fear in human amygdala. Nature Neuroscience, 19(8), 1041–1049.

Michelmann, S., Bowman, H., & Hanslmayr, S. (2016). The Temporal Signature of Memories: Identification of a General Mechanism for Dynamic Memory Replay in Humans. PLOS Biology, 14(8), e1002528. 10.1371/journal.pbio.1002528

Milad, M. R., & Quirk, G. J. (2012). Fear extinction as a model for translational neuroscience: Ten years of progress. Annual Review of Psychology, 63(1), 129–151.

Miller, J. F., Neufang, M., Solway, A., Brandt, A., Trippel, M., Mader, I., Hefft, S., Merkow, M., Polyn, S. M., Jacobs, J., Kahana, M. J., & Schulze-Bonhage, A. (2013). Neural Activity in Human Hippocampal Formation Reveals the Spatial Context of Retrieved Memories. Science, 342(6162), 1111–1114. 10.1126/science.1244056

Mueller, E. M., Panitz, C., Hermann, C., & Pizzagalli, D. A. (2014). Prefrontal Oscillations during Recall of Conditioned and Extinguished Fear in Humans. The Journal of Neuroscience, 34(21), 7059. 10.1523/JNEUROSCI.3427-13.2014

Neumann, D. L., & Kitlertsirivatana, E. (2010). Exposure to a novel context after extinction causes a renewal of extinguished conditioned responses: Implications for the treatment of fear. Behaviour Research and Therapy, 48(6), 565–570. 10.1016/j.brat.2010.03.002

Neumann, D. L., Lipp, O. V., & Cory, S. E. (2007). Conducting extinction in multiple contexts does not necessarily attenuate the renewal of shock expectancy in a fear-conditioning procedure with humans. Behaviour Research and Therapy, 45(2), 385–394. 10.1016/j.brat.2006.02.001

Nguyen, R., Koukoutselos, K., Forro, T., & Ciocchi, S. (2023). Fear extinction relies on ventral hippocampal safety codes shaped by the amygdala. Science Advances, 9(22), eadg4881. 10.1126/sciadv.adg4881

Oostenveld, R., Fries, P., Maris, E., & Schoffelen, J.-M. (2011). FieldTrip: Open source software for advanced analysis of MEG, EEG, and invasive electrophysiological data. Computational Intelligence and Neuroscience, 2011, 156869. 10.1155/2011/156869

Orsini, C. A., Kim, J. H., Knapska, E., & Maren, S. (2011). Hippocampal and prefrontal projections to the basal amygdala mediate contextual regulation of fear after extinction. Journal of Neuroscience, 31(47), 17269–17277.

Orsini, C. A., & Maren, S. (2012). Neural and cellular mechanisms of fear and extinction memory formation. Memory Formation, 36(7), 1773–1802. 10.1016/j.neubiorev.2011.12.014

Pacheco Estefan, D. (2023). How Can We Track Cognitive Representations with Deep Neural Networks and Intracranial EEG? In N. Axmacher (Ed.), Intracranial EEG: A Guide for Cognitive Neuroscientists (pp. 849–862). Springer International Publishing. 10.1007/978-3-031-20910-9_50

Pacheco Estefan, D., Sánchez-Fibla, M., Duff, A., Principe, A., Rocamora, R., Zhang, H., Axmacher, N., & Verschure, P. F. M. J. (2019). Coordinated representational reinstatement in the human hippocampus and lateral temporal cortex during episodic memory retrieval. Nature Communications, 10(1), 1–13.

Pacheco Estefan, D., Zucca, R., Arsiwalla, X., Principe, A., Zhang, H., Rocamora, R., Axmacher, N., & Verschure, P. F. M. J. (2021). Volitional learning promotes theta phase coding in the human hippocampus. Proceedings of the National Academy of Sciences, 118(10).

Pacheco-Estefan, D., Fellner, M.-C., Kunz, L., Zhang, H., Reinacher, P., Roy, C., Brandt, A., Schulze-Bonhage, A., Yang, L., Wang, S., Liu, J., Xue, G., & Axmacher, N. (2024). Maintenance and transformation of representational formats during working memory prioritization. Nature Communications, 15(1), 8234. 10.1038/s41467-024-52541-w

Paré, D., & Collins, D. R. (2000). Neuronal Correlates of Fear in the Lateral Amygdala: Multiple Extracellular Recordings in Conscious Cats. The Journal of Neuroscience, 20(7), 2701. 10.1523/JNEUROSCI.20-07-02701.2000

Parvizi, J., & Kastner, S. (2018). Promises and limitations of human intracranial electroencephalography. Nature Neuroscience, 21(4), 474–483.

Pirazzini, G., Starita, F., Ricci, G., Garofalo, S., di Pellegrino, G., Magosso, E., & Ursino, M. (2023). Changes in brain rhythms and connectivity tracking fear acquisition and reversal. Brain Structure and Function, 228(5), 1259–1281. 10.1007/s00429-023-02646-7

Popescu, A. T., Popa, D., & Paré, D. (2009). Coherent gamma oscillations couple the amygdala and striatum during learning. Nature Neuroscience, 12(6), 801–807. 10.1038/nn.2305

Qasim, S. E., Mohan, U. R., Stein, J. M., & Jacobs, J. (2023). Neuronal activity in the human amygdala and hippocampus enhances emotional memory encoding. Nature Human Behaviour, 7(5), 754–764. 10.1038/s41562-022-01502-8

Quinn, J. J., Ma, Q. D., Tinsley, M. R., Koch, C., & Fanselow, M. S. (2008). Inverse temporal contributions of the dorsal hippocampus and medial prefrontal cortex to the expression of long-term fear memories. Learning & Memory, 15(5), 368–372.

Quirk, G. J., & Mueller, D. (2008). Neural Mechanisms of Extinction Learning and Retrieval. Neuropsychopharmacology, 33(1), 56–72. 10.1038/sj.npp.1301555

Rahman, M. M., Shukla, A., & Chattarji, S. (2018). Extinction recall of fear memories formed before stress is not affected despite higher theta activity in the amygdala. eLife, 7, e35450. 10.7554/eLife.35450

Rescorla, R. A., & Wagner, A. R. (1972). A theory of Pavlovian conditioning: Variations in the effectiveness of reinforcement and non-reinforcement. *Classical Conditioning*, Current Research and Theory, 2, 64–69.

Rivière, D., Geffroy, D., Denghien, I., Souedet, N., & Cointepas, Y. (2011). Anatomist: A python framework for interactive 3D visualization of neuroimaging data. 3–4.

Rolls, E. T. (1999). Spatial view cells and the representation of place in the primate hippocampus. Hippocampus, 9(4), 467–480.

Saint Amour di Chanaz, L., Pérez-Bellido, A., Wu, X., Lozano-Soldevilla, D., Pacheco-Estefan, D., Lehongre, K., Conde-Blanco, E., Roldan, P., Adam, C., Lambrecq, V., Frazzini, V., Donaire, A., Carreño, M., Navarro, V., Valero-Cabré, A., & Fuentemilla, L. (2023). Gamma amplitude is coupled to opposed hippocampal theta-phase states during the encoding and retrieval of episodic memories in humans. Current Biology, 33(9), 1836–1843.e6. 10.1016/j.cub.2023.03.073

Santini, E., Quirk, G. J., & Porter, J. T. (2008). Fear Conditioning and Extinction Differentially Modify the Intrinsic Excitability of Infralimbic Neurons. The Journal of Neuroscience, 28(15), 4028. 10.1523/JNEUROSCI.2623-07.2008

Seger, S. E., Kriegel, J. L. S., Lega, B. C., & Ekstrom, A. D. (2023). Memory-related processing is the primary driver of human hippocampal theta oscillations. Neuron, 111(19), 3119–3130.e4. 10.1016/j.neuron.2023.06.015

Seidenbecher, T., Laxmi, T. R., Stork, O., & Pape, H.-C. (2003). Amygdalar and Hippocampal Theta Rhythm Synchronization During Fear Memory Retrieval. Science, 301(5634), 846–850. 10.1126/science.1085818

Shiban, Y., Pauli, P., & Mühlberger, A. (2013). Effect of multiple context exposure on renewal in spider phobia. Behaviour Research and Therapy, 51(2), 68–74. 10.1016/j.brat.2012.10.007

Sperl, M. F. J., Panitz, C., Rosso, I. M., Dillon, D. G., Kumar, P., Hermann, A., Whitton, A. E., Hermann, C., Pizzagalli, D. A., & Mueller, E. M. (2019). Fear Extinction Recall Modulates Human Frontomedial Theta and Amygdala Activity. Cerebral Cortex, 29(2), 701–715. 10.1093/cercor/bhx353

Staresina, B. P., Bergmann, T. O., Bonnefond, M., van der Meij, R., Jensen, O., Deuker, L., Elger, C. E., Axmacher, N., & Fell, J. (2015). Hierarchical nesting of slow oscillations, spindles and ripples in the human hippocampus during sleep. Nature Neuroscience, 18(11), 1679–1686. 10.1038/nn.4119

Staresina, B. P., Michelmann, S., Bonnefond, M., Jensen, O., Axmacher, N., & Fell, J. (2016). Hippocampal pattern completion is linked to gamma power increases and alpha power decreases during recollection. eLife, 5, e17397. 10.7554/eLife.17397

Stujenske, J. M., Likhtik, E., Topiwala, M. A., & Gordon, J. A. (2014). Fear and Safety Engage Competing Patterns of Theta-Gamma Coupling in the Basolateral Amygdala. Neuron, 83(4), 919–933. 10.1016/j.neuron.2014.07.026

Szeska, C., Richter, J., Wendt, J., Weymar, M., & Hamm, A. O. (2020). Promoting long-term inhibition of human fear responses by non-invasive transcutaneous vagus nerve stimulation during extinction training. Scientific Reports, 10(1), 1529. 10.1038/s41598-020-58412-w

Taschereau-Dumouchel, V., Kawato, M., & Lau, H. (2020). Multivoxel pattern analysis reveals dissociations between subjective fear and its physiological correlates. Molecular Psychiatry, 25(10), 2342–2354. 10.1038/s41380-019-0520-3

Taub, A. H., Perets, R., Kahana, E., & Paz, R. (2018). Oscillations synchronize amygdala-to-prefrontal primate circuits during aversive learning. Neuron, 97(2), 291–298.

Tulving, E. (2002). Episodic memory: From mind to brain. Annual Review of Psychology, 53(1), 1–25.

Visser, R. M., Scholte, H. S., Beemsterboer, T., & Kindt, M. (2013). Neural pattern similarity predicts long-term fear memory. Nature Neuroscience, 16(4), 388–390. 10.1038/nn.3345

Wang, Y., Olsson, S., Lipp, O. V., & Ney, L. J. (2024). Renewal in human fear conditioning: A systematic review and meta-analysis. Neuroscience & Biobehavioral Reviews, 159, 105606. 10.1016/j.neubiorev.2024.105606

Weber, J., Iwama, G., Solbakk, A.-K., Blenkmann, A. O., Larsson, P. G., Ivanovic, J., Knight, R. T., Endestad, T., & Helfrich, R. (2023). Subspace partitioning in the human prefrontal cortex resolves cognitive interference. Proceedings of the National Academy of Sciences, 120(28), e2220523120. 10.1073/pnas.2220523120

Wu, X., Packard, P. A., García-Arch, J., Bunzeck, N., & Fuentemilla, L. (2023). Contextual incongruency triggers memory reinstatement and the disruption of neural stability. NeuroImage, 273, 120114. 10.1016/j.neuroimage.2023.120114

Xie, W., Wittig, J. H., & Zaghloul, K. A. (2023). What Do I Need to Consider for Multivariate Analysis of iEEG Data? In N. Axmacher (Ed.), Intracranial EEG: A Guide for Cognitive Neuroscientists (pp. 557–566). Springer International Publishing. 10.1007/978-3-031-20910-9_34

Xue, G., Dong, Q., Chen, C., Lu, Z., Mumford, J. A., & Poldrack, R. A. (2010). Greater Neural Pattern Similarity Across Repetitions Is Associated with Better Memory. Science, 330(6000), 97–101. 10.1126/science.1193125

Yaffe, R. B., Kerr, M. S. D., Damera, S., Sarma, S. V., Inati, S. K., & Zaghloul, K. A. (2014). Reinstatement of distributed cortical oscillations occurs with precise spatiotemporal dynamics during successful memory retrieval. Proceedings of the National Academy of Sciences, 111(52), 18727–18732. 10.1073/pnas.1417017112

Yonelinas, A. P., Ranganath, C., Ekstrom, A. D., & Wiltgen, B. J. (2019). A contextual binding theory of episodic memory: Systems consolidation reconsidered. Nature Reviews Neuroscience, 20(6), 364–375. 10.1038/s41583-019-0150-4

Zhang, H., Skelin, I., Ma, S., Paff, M., Mnatsakanyan, L., Yassa, M. A., Knight, R. T., & Lin, J. J. (2024). Awake ripples enhance emotional memory encoding in the human brain. Nature Communications, 15(1), 215. 10.1038/s41467-023-44295-8

Zhang, X., Kim, J., & Tonegawa, S. (2020). Amygdala Reward Neurons Form and Store Fear Extinction Memory. Neuron, 105(6), 1077–1093.e7. 10.1016/j.neuron.2019.12.025

Zheng, J., Anderson, K. L., Leal, S. L., Shestyuk, A., Gulsen, G., Mnatsakanyan, L., Vadera, S., Hsu, F. P. K., Yassa, M. A., Knight, R. T., & Lin, J. J. (2017). Amygdala-hippocampal dynamics during salient information processing. Nature Communications, 8(1), 14413. 10.1038/ncomms14413

