## Supplementary Material for "Representational dynamics during extinction of fear memories in the human brain"

Pacheco-Estefan et al., 2025

Supplementary Notes 1-6

Supplementary Figures 1-3

Supplementary Tables 1-3

Supplementary Video 1

Supplementary References

**Supplementary Note 1. No differences in coordination of item stability and context specificity across cue types.**

We computed item stability and context specificity separately for each of our cue types (CS++, CS+- and CS--), focusing on the ROIs where we observed significant effects in the analysis including all trials during extinction. We then assessed whether coordination differed between CS+ vs CS- trials, and between CS++ vs CS+- trials. We performed the analysis across all time points in the temporal generalization map and assessed statistical significance via cluster-based permutation statistics, shuffling the condition labels 1,000 times. We only considered significant those time-by-time clusters whose summed t-values ranked above the 95th percentile of the null distribution.

In most analyses, we did not observe large clusters in the temporal generalization maps, as observed in the resulting p-values presented in the table below (corresponding to the rank of these clusters within the distribution of surrogate cluster). Thus, while our main results indicate that item stability differed between experimental conditions in the TMP and the AMY (Figure 3), the coordination of item stability across brain regions did not differ between conditions.

| Analysis | Trial  Region | CS+ vs CS- | CS++ vs CS+- |
| --- | --- | --- | --- |
| Item stability | TMP-AMY | N= 22, p = 0.69 | N = 19, p = 0.69 |
|  | TMP-HPC | N= 23, p = 0.8 | N = 21, p = 0.38 |
|  | TMP-lPFC | N= 17, p = 0.51 | N = 16, p = 0.81 |
|  | TPM-OFC | N= 9, p = 0.59 | N = 9, p = 0.93 |
|  | AMY-HPC | N= 24, p = 0.34 | N = 23, p = 0.86 |
| Context specificity | lPFC-AMY | N = 14, p = 0.91 | N = 14, p = 0.95 |
|  | lPFC-TMP | N= 17, p = 0.21 | N = 16, p = 0.64 |

Summary table presents the results of the contrasts between CS+ and CS- trials, as well as between CS++ and CS+- trials, in the analyses of the coordination of both item stability and context specificity between regions. Analyses were conducted at all time points in the temporal generalization maps, and the reported p-values reflect the rank of the largest observed cluster in relation to a null distribution constructed by shuffling the condition labels. The number of participants (N) included in each analysis is also indicated.

**Supplementary Note 2. No differences in correlations of item stability and context specificity with AMY_THETA_ across cue types.**

To assess differences in the correlations between AMY_THETA_ and item stability or context specificity across conditions, we extracted a single-trial metric of AMY_THETA_ based on the cluster of significant differences between CS+ and CS- items observed during extinction (Figure 2; see as schematic depiction in Figure 5A, left). We separately z-scored this metric for CS+ and CS- trials, in order to avoid any spurious correlation of single-trial values driven by main condition differences. In addition, we averaged item stability and context specificity across time in the time period of AMY_THETA_ effects in each trial and each ROI. We correlated AMY theta power with item stability and context specificity across trials, separately in our three trial types (CS++, CS+- and CS--), and assessed condition differences using a one-way repeated measures ANOVA.

Our results did not reveal any significant difference between conditions in the AMY (F(2, 44) = 1.15, p = 0.32). Post-hoc tests revealed that none of the pairwise comparisons showed differences, even at an uncorrected level (all p > 0.19). Similarly, in the HPC, correlations did not differ between conditions (F(2, 34) = 0.2, p = 0.82) and pairwise correlations were not significantly different (all p > 0.49). Finally, the analysis of lPFC context specificity revealed no significant differences between conditions (F(2, 22) = 0.91, p = 0.42), and no significant differences in the pairwise post hoc tests (all p > 0.2).

These results demonstrate that the observed trial-level correlations between AMY_THETA_ and item stability in AMY and HPC (Figure 5) did not differ across trial types. In addition, they show that the correlations between AMY_THETA_ and lPFC_CONTEXT_ are not significantly different across conditions. Together, these findings suggest that AMY theta oscillations play a role in the representation of cues and contexts throughout the extinction network irrespective of their valence.

**Supplementary Note 3. Reinstatement analyses across cue types**

We compared the reinstatement values between our three trial types (CS++, CS+-, CS--), separately for acquisition-to-test and extinction-to-test reinstatement, in the brain regions where we observed significantly higher item-stability for the CS+ as compared to the CS- trials during extinction, i.e., the AMY and the TMP. In addition, we compared the differential reinstatement of memory traces (acquisition-to-test minus extinction-to-test) across these conditions. In a separate analysis, we investigated whether the amount of acquisition-to-test reinstatement differed from the amount of extinction-to-test reinstatement for any of the categories.

For the first analysis, we assessed possible condition effects on reinstatement during time windows of 500ms during acquisition/extinction and test, sliding in 50ms (90% overlap). Only matching time-windows were considered for this analysis because a temporal generalization analysis would not be possible for the differential reinstatement, which involves time periods during three phases (acquisition, extinction, and test). We averaged across trials and across time within each 500ms window for each participant and performed one-way repeated-measures ANOVAs at each time bin to assess statistical significance. In the AMY, we did not observe any significant differences between our three conditions in the acquisition-to-test reinstatement (all time bins: F < 1.73, p > 0.18), the extinction-to-test reinstatement (all time bins: F < 2.8, all p > 0.06) or in the differential reinstatement analysis (all time bins: F < 2.54, all p > 0.08), even before applying correction for multiple comparisons. Similarly, in the TMP, we found no significant condition differences, even at an uncorrected level, for acquisition-to-test reinstatement (all time bins: F < 1.4, p > 0.26), extinction-to-test reinstatement (all time bins: F < 2.53, p > 0.089) or in the differential reinstatement analysis (all time bins: F < 2.42, p > 0.09; see Figure below). These results suggest that reinstatement does not differ between conditions.

In the second analysis, we compared the magnitude of acquisition-to-test and extinction-to-test reinstatement separately for the CS++, the CS+- and the CS-- trials. We focused on the time periods where significant item-stability effects were observed in each region (AMY: 1.25-1.5s; TMP: 0.65-1s). In the TMP, none of the cue types showed significant differences between acquisition-to-test and extinction-to-test reinstatement (CS++: t(25)= -1.11; p = 0.28; CS+-: t(25)= 0.647; p = 0.52; CS--: t(25)= 1.59; p = 0.12). In the AMY, we observed no significant differences in CS++ (t(30)= -0.092; p = 0.93) or CS-- trials (t(30)= -0.352; p = 0.728). However, for the CS+- trials, we observed a trend toward higher levels of acquisition-to-test reinstatement compared to extinction-to-test reinstatement (t(30)= 1.72; p = 0.096).

**Supplementary Note 4: Paradigm main features and rationale**

The experimental paradigm we conducted is novel and its unique combination of features has not been validated in previous studies. However, each of the design decisions we made was based on practical and theoretical considerations derived from the fear conditioning literature. We below present a rationale for each of these design decisions.

First, we employed a multi-cue conditioning protocol in which one stimulus (CS+) predicts the unconditioned stimulus (US), while another (CS−) does not. This approach is the most commonly employed in human fear conditioning studies and differs from paradigms in which a single cue is first associated with a US and later extinguished (Lonsdorf et al., 2017).

Second, like previous research investigating the representational dynamics of fear learning (Visser et al., 2013), we used natural images as conditioned stimuli rather than auditory stimuli—more employed in animal research—or simplified shapes (Chen et al., 2021; Neumann et al., 2007; Schmitz & Grillon, 2012). Our approach is not only arguably more ecologically valid but also better suited to capture cue representations across several sensory areas including association cortices.

Third, our study follows an ABC design, in which acquisition occurs in one context, extinction in another, and testing in a novel third context. This structure is commonly employed in both rodent and human studies that examine the role of context in fear extinction (Balooch et al., 2012; Corcoran & Maren, 2001, 2004; Effting & Kindt, 2007; Hermann et al., 2016; Milad et al., 2005). However, it differs from other human studies that have used ABA (Krisch et al., 2018) or ABB (Balooch et al., 2012) paradigms to investigate context-dependent fear extinction. We employed an ABC paradigm because we aimed to investigate whether the re-occurrence of fear memories during the final test phase was determined by the similarities of internal (i.e., neural) representations of contexts during this phase to context representations during either the acquisition or the extinction phase. This differs from ABA and ABB paradigms in which the presented contexts during the test phase are either identical to the contexts during acquisition (ABA) or during extinction (ABB).

Notably, while our paradigm follows an ABC structure, our design includes the presentation of several distinct yet thematically related context videos in each experimental phase (e.g., videos of sea landscapes). This was done to facilitate the analysis of context-specific representations using Representational Similarity Analysis (RSA), which requires the presentation of multiple instances of a particular context. In particular, as stated in our hypothesis, we aimed to evaluate the generalization of context representations during acquisition and their distinctiveness during extinction.

Fourth, we applied a partial reinforcement rate of 50% (Dunsmoor et al., 2007, 2012, 2014; Visser et al., 2013). While some studies have used slightly higher rates, such as 60% (Battaglia et al., 2018; Milad et al., 2007), 62.5% (Hermann et al., 2016), or 66% (Dunsmoor & Murphy, 2014), we opted for 50% to promote a more gradual acquisition and extinction of fear associations (note that both processes are affected by this choice). Slower rates also reduce the risk of habituation to the unconditioned stimulus (US), which was particularly important in our study given the relatively mild US we employed (see next point).

Fifth, in our paradigm, the US consisted of a human female face with a neutral expression following CS− items and a fearful expression following (50% of) CS+ items. Presentation of the fearful face following CS+ items was, in addition, paired with a loud scream. The choice of using emotional facial expressions was informed by studies suggesting that negative facial expressions increase resistance to extinction compared to neutral faces (Mineka & Öhman, 2002; Ney et al., 2022; Öhman & Öst, 1985). Similarly, pairing this face with a scream was grounded on previous fear conditioning research in sensitive populations such as infants and patients (Glenn, Klein, et al., 2012; Hamm et al., 1989). Moreover, the face-scream protocol has been shown to produce similar—though milder—responses in fear-potentiated startle and self-reported fear ratings in healthy participants as compared to electric shocks (Glenn, Lieberman, et al., 2012).

Sixth, to strengthen the association between the CS and US, we incorporated a narrative element, or "cognitive embedding". Previous research has shown that semantically or perceptually meaningful CS-US pairings are learned more effectively and are more resistant to extinction than arbitrary associations (Garcia & Koelling, 1966; Hamm et al., 1989). In our paradigm, we introduced the character of Nina, a backpacker who travels the world and stays in hotels where electric devices occasionally malfunction. Pictures of these electric devices were used as our CS (a fan, a hairdryer and a toaster). The negative emotional expression of “Nina’s” face, paired with the scream, aligns with this narrative.

Seventh, we used videos to represent contextual information rather than simpler stimuli such as colored backgrounds (Kalisch et al., 2006), images of scenes or rooms (Battaglia et al., 2018; Xia et al., 2023), or more complex VR environments (Baas et al., 2004; Glotzbach-Schoon et al., 2013). This was made to enhance realism and reinforce the narrative of a traveler visiting diverse regions with distinct landscapes and climates while avoiding the difficulties of conducting VR research in clinical populations. In addition, we used videos since previous research has shown that their neural representations can be particularly well tracked in human electrophysiological data (Michelmann et al., 2016).

Eighth, as in various human fear conditioning studies that rely on self-reports rather than physiological measures, we assessed participants' expectancy ratings of safety and threat rather than collecting electrodermal activity data (Bandarian Balooch & Neumann, 2011; Boddez et al., 2013). This was guided by both ethical and practical considerations, particularly given the constraints of conducting research with implanted epilepsy patients, and to emphasize the role of conscious awareness in learning and extinction.

Ninth, our testing phase occurred shortly after extinction without introducing additional delays. This decision was driven by the limited availability of patients in the clinical context. We acknowledge that some studies have implemented delays between extinction and testing to evaluate the stability of extinction in the long-term (Milad et al., 2005; Xia et al., 2023), but also highlight that several others have applied immediate testing (e.g., (Vansteenwegen et al., 2007; Visser et al., 2013).

Finally, due to the time constraints of conducting research with implanted patients, we did not conduct individualized aversiveness testing for the US. Instead, as in previous studies (Schmitz & Grillon, 2012; Wehrli et al., 2022), we selected a notoriously aversive stimulus—a loud scream—and adjusted its duration and intensity to a highly unpleasant level (1s, ~85dB).

The unique and novel combination of features in our paradigm—including the use of multiple, thematically related videos as contextual stimuli, the ABC structure, the 50% reinforcement rate, and the embedding of the experiment into a plausible narrative—has not been validated in previous research. However, as we have discussed, these methodological choices were carefully informed by the fear conditioning literature in humans. Notably, a very similar version of this paradigm, developed by our group for an fMRI study, includes the same stimuli, a very similar overall structure, and the same cover story but was conducted in healthy participants and employed a different US (electric shocks). This study has been recently published in biorXiv and is currently under review (Bouyeure et al., 2024). While these data are beyond the current study, they provide an additional validation of our core paradigm choices

**Supplementary Note 5: Different electrode configurations did not affect volume of neural tissue sampled and main results**

In our study, data were collected using three types of electrodes: two different types in Paris (micro-macro: 21 electrodes; macro-only: 176 electrodes) and one from Guangzhou (434 electrodes). These electrodes have variable surface area and inter-contact distances (Paris micro-macro: 1.6/3-7mm; Paris macro-only: 1.7/5mm; Guangzhou: 2/3.5mm). Additionally, within a single micro-macro electrode in Paris, the first and second contacts are separated by 3 mm, while subsequent contacts (e.g., 2 to 3, 3 to 4) are spaced 7 mm apart.

Differences in electrode configurations, combined with our use of a bipolar referencing scheme, might have affected the total volume of neural tissue sampled in our two patient groups in Paris and Guangzhou. However, it is unlikely that such differences affected our main results for several reasons. First, we only included electrodes located in gray matter, which removed a large number of contact points within each electrode (~50%). The removal of many electrode contacts introduced variability in sampling density, affecting both patient cohorts similarly. Second, a higher sampling density does not necessarily equate to more information, as closely spaced electrodes in iEEG often capture activity from the same sources and exhibit similar neurophysiological profiles. We note that most of our analyses involved averaging electrode activity in each subject independently before performing statistical comparisons at the group level, which further reduces the risk that sampling density differences systematically biased our findings. Third, many of our key findings were observed in the AMY, which had the lowest sampling density overall, with a mean number of electrodes per subject of 2.56 ± 1.39. Notably, 10 out of 22 patients had only a single electrode in the AMY, making it unlikely that inter-electrode distances or contact surface area significantly influenced our results in this region. Fourth, the inter-contact spacing of the first two electrodes in the micro-macro arrays from Paris (3 mm) closely resembles that of the electrode configurations used in Guangzhou (3.5 mm), minimizing potential sampling density differences in these electrodes.

We performed two control analyses to corroborate that different electrode configurations in our patient’s cohort did not affect the density of neural tissue sampled. First, we computed the Delaunay triangulation of electrode configurations, a method that quantifies electrode density by forming triangles between contact points and calculating the mean triangle area. Focusing on our largest and more densely implanted ROI (the TMP) we evaluated triangulation in the left and right hemispheres in each subject independently. Only subjects with at least three electrodes in the TMP (N=36) were included in this analysis, as a minimum of three electrodes is required to form triangles. Statistical significance was assessed using a two-way ANOVA with "Recording site" (Guangzhou/Paris) and "Hemisphere" as factors. Results showed no significant difference in mean triangle area between hemispheres (F(1,32) = 2.61, p = 0.11) or between patient cohorts (F(1,32) = 0.96, p = 0.33). These findings indicate that although electrode spacing varied due to differences in electrode configurations, the overall sampling density remained consistent across our two patient groups.

To further corroborate that the different electrode configurations did not affect our results, we conducted our oscillatory power analyses in the AMY separately for Guangzhou and Paris patients. We specifically focused on the significant cluster observed in the main analysis contrasting “current” valence (CS+ vs CS-) during extinction. We observed the same pattern of results across our two cohorts of patients, with higher theta power for CS- as compared to CS+ items during extinction (Guangzhou: t(19)= -4.32, p = 0.0004; Paris: t(11)= 2.93, p = 0.014). Taken together, the results of these analyses suggest that our two patient cohorts did not show prominent differences in the neural tissue sampled in the TMP, and that the AMY theta power results during extinction were consistent in our two patient groups (see Supplementary Figure 2).

Finally, we highlight that recent years have seen an increase of multi-center iEEG research, which involves collecting and analyzing data from multiple epilepsy centers to generate larger and more diverse datasets. Several studies have successfully integrated multi-center iEEG data despite variations in electrode configurations, including even combination of subdural strips and grids with depth electrodes, which differ in regard to sampling density even more than different types of depth electrodes do (Bernabei et al., 2023; Dimakopoulos et al., 2022; Henin et al., 2021). While there is currently no established consensus or guideline on how to address sampling differences in the literature, we believe this is an important topic of research in the future, as multi-center iEEG studies are becoming more prominent.

**Supplementary Note 6: Connectivity analysis using the weighted phase lag index**

Conceptually, we were predominantly interested in investigating the representations of contexts and items during fear learning and extinction. For this reason, we employed metrics of representational coordination rather than standard connectivity analyses in order to investigate the propagation of representational signatures throughout the fear and extinction network. However, we acknowledge that connectivity metrics could offer complementary insights into the neural mechanisms of fear extinction.

Over the past decades, several connectivity metrics have been developed and applied. One of the first to be proposed was the Phase Locking Value (PLV) (Lachaux et al., 1999), which quantifies the consistency of phase angle differences between two signals. The PLV has been widely applied in cognitive neuroscience (Mormann et al., 2000; Pacheco Estefan et al., 2019; Varela et al., 2001), but this metric is highly susceptible to volume conduction since it includes apparent connectivity at phase differences at zero lag. Volume conduction refers to the phenomenon where electrical activity from neuronal sources propagates almost instantaneously through brain tissues, leading to signal spread. This effect can artificially inflate metrics of connectivity in iEEG, as different electrodes might capture activity from the same neurophysiological source.

To circumvent the problem of volume conduction, several metrics based on phase lag were developed in the last years. In particular, phase-based connectivity measures were established that ignore zero phase-lag connectivity, including the phase-slope index (Nolte et al., 2008), the phase lag index (Stam et al., 2007), and the weighted phase-lag index (Vinck et al., 2011). The Phase Lag Index (PLI) quantifies asymmetries in phase lead/lag, ignoring synchronized activity at zero or 180° phase lag—which is often the result of volume conduction rather than true connectivity. The weighted PLI (wPLI) assigns greater weight to larger phase differences by considering the magnitude of the imaginary component of the cross-spectrum, effectively suppressing small phase differences that may result from volume conduction. By improving robustness against noise as compared to the PLI and minimizing false connectivity, wPLI has become a widely employed metric for mapping functional networks in EEG/iEEG research (Imperatori et al., 2019; Vinck et al., 2011).

To complement our representational coordination analyses and provide a more comprehensive description of how fear extinction signals propagate throughout the fear and extinction network, we assessed the wPLI between pairs of ROIs. We specifically focused on the ROIs where we observed significant coordination in our item stability (Figure 3D) or context specificity (Figure 4D) analyses. These included AMY-TMP, AMY-HPC, AMY-lPFC, TMP-HPC, TMP-OFC, and TMP-lPFC. We computed wPLI across trials for each pair of electrodes in their correspondent ROI pairs, focusing on the theta (4–8 Hz) frequency band, as this was the frequency range where we observed the most prominent effects in the AMY in our data. Notably, phase-based connectivity in the theta range has been associated with long-range neural communication (Fell & Axmacher, 2011). We specifically selected the time periods where we observed significant coordination effects in the original analysis (see table below).

We computed wPLI for all electrode pairs within predefined ROI pairs, separately for each subject and each of our three cue types (CS++, CS+-, and CS--). In subjects with bi-hemispheric implants, we computed wPLI for channel pairs within the same hemisphere only. wPLI metrics obtained for each electrode pair were averaged across electrodes in each subject before performing statistical comparisons. Significant differences were assessed with a repeated measures one-way ANOVA.

Our results did not reveal any significant differences in wPLI during the time periods of significant coordination in any of the ROI pairs even at an uncorrected level (see table below).

| **ROIs** | **Time period** | **ANOVA results** |
| --- | --- | --- |
| AMY-TMP | 0-1.75s | F(2,24) = 2.22, p = 0.12 |
| AMY-HPC | 0.5-1.75s | F(2,20) = 1.08, p = 0.34 |
| AMY-lPFC | 1-1.65s | F(2,10) = 0.16, p = 0.85 |
| TMP-HPC | 0-1.45 | F(2,22) = 0.09, p = 0.91 |
| TMP-OFC | 0.15-1s | F(2,7) = 0.72, p = 0.5 |
| TMP-lPFC | 0-0.8s | F(2,14) = 0.24, p = 0.78 |

The absence of significant differences in our phase-based connectivity analysis suggests that the observed coordination in representational signals in our study is not linked to phase-based connectivity in the theta frequency range. However, we acknowledge that the space of possible connectivity analyses is vast, and our metric of representational coordination may align more closely with other forms of connectivity, such as power-based or causal connectivity, which could be investigated more comprehensively in the future.

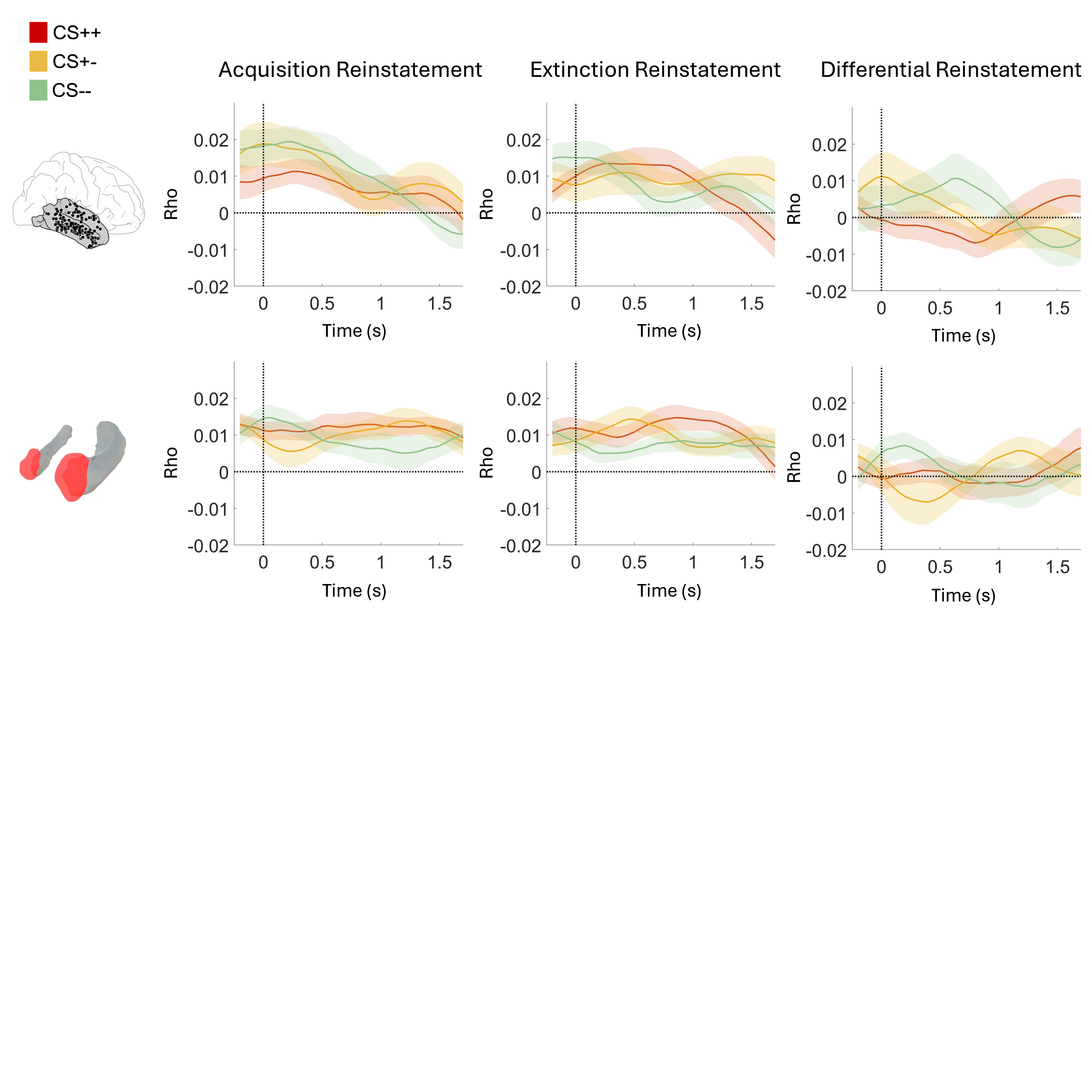

**Supplementary Figure 1. Reinstatement of acquisition and extinction memory traces do not differ across trial types**. Acquisition-to-test (left), extinction-to-test and differential reinstatement (acquisition minus extinction-to-test) did not differ between our three trial types (CS++, CS+-, CS--).

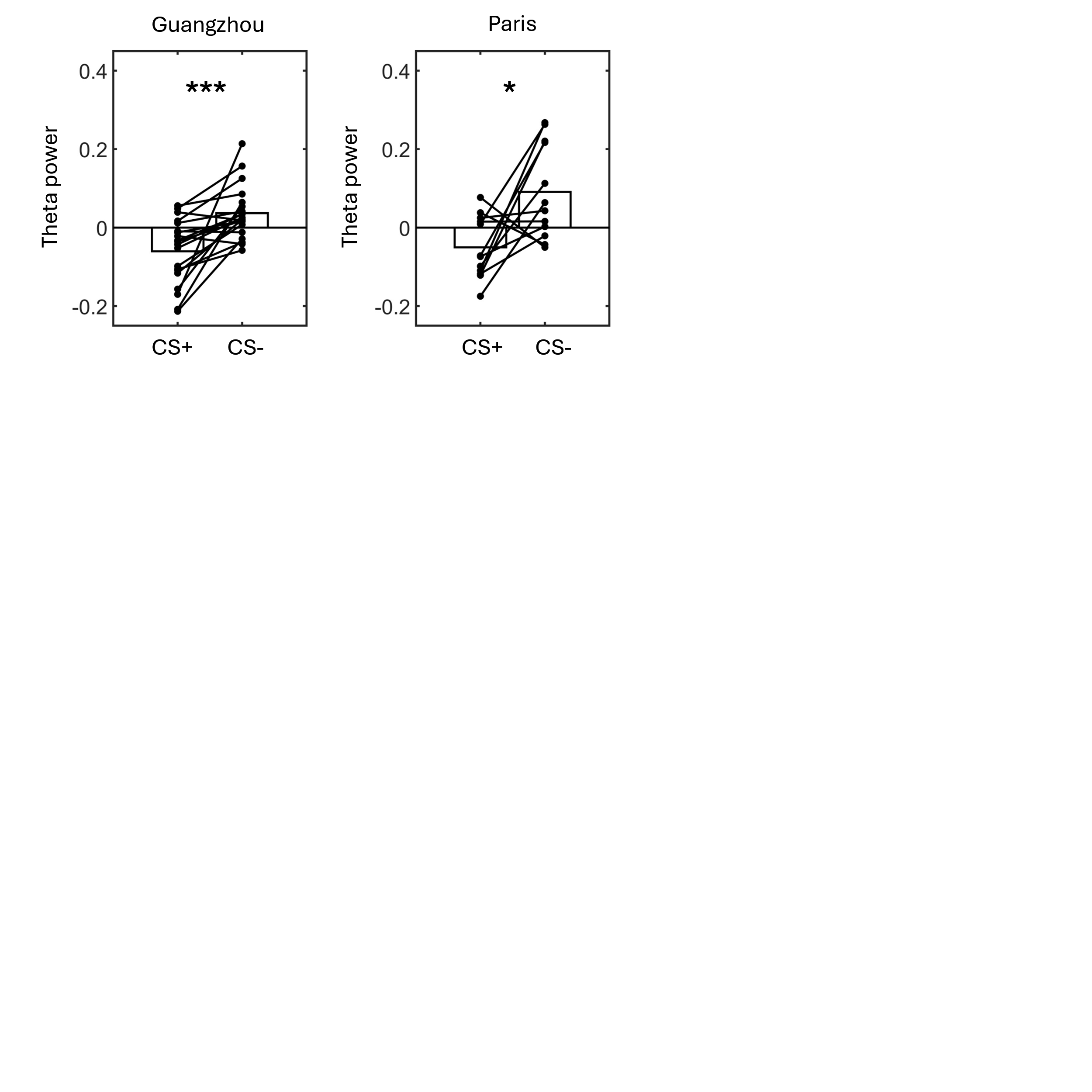
**Supplementary Figure 2. Theta power in the AMY during extinction was significantly higher for CS- as compared to CS+ trials in both the Guangzhou and the Paris patient cohorts.**

Figure shows mean AMY power within the time-frequency cluster identified in the main analysis during extinction (Figure 2A, bottom left) for the Guangzhou (left) and the Paris (right) patient groups. In both cohorts, AMY theta power was significantly higher for CS- as compared to CS+ trials, consistent with the findings observed when combining the data of the two groups. These results demonstrate that our AMY theta power findings are robust and not influenced by methodological differences between the epilepsy centers where the data was collected (e.g., variations in electrode configurations or recording devices in Paris and Guangzhou).

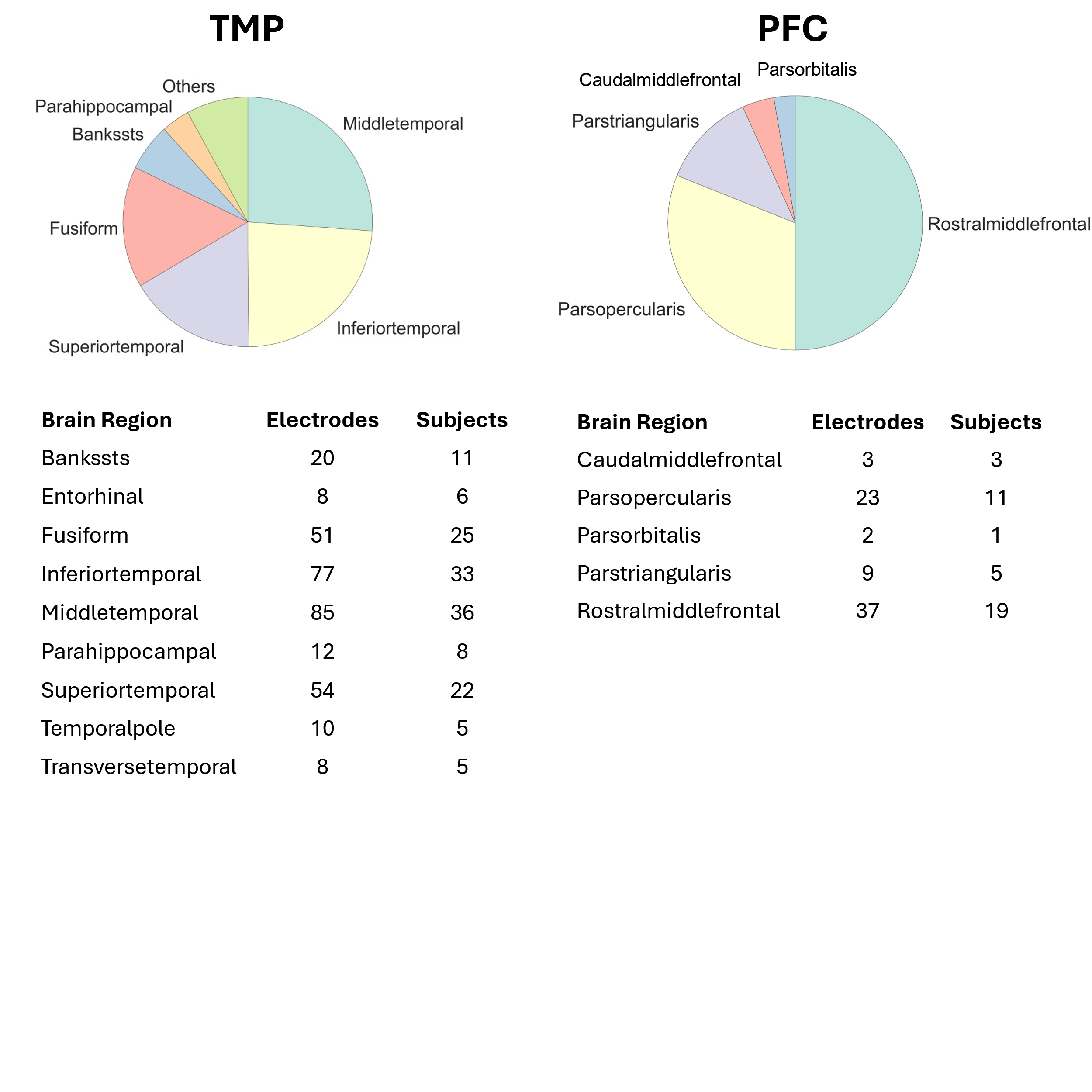

**Supplementary Figure 3. Proportion of electrodes in TMP and PFC subregions.**

Proportion of electrodes included in specific subregions of the TMP (left) and the PFC (right; Desikan Killian atlas; Desikan et al., 2006) across all subjects. A summary table for each ROI, including the number of electrodes and the number of subjects in each subregion, is presented below the pie plots.

| **Participant** | **Gender** | **Age** | **Global IQ** | **Seizure Onset Zone** | **MQ** |
| --- | --- | --- | --- | --- | --- |
| DBX | M | 22 | 115 | Right temporal lobe, bilateral medial temporal lobes, insula, and right extra-orbital cortex | 106 |
| LSY | F | 16 | 88 | Right medial temporal lobe | 39 |
| JJL | F | 25 | 86 | Left temporo-occipital lobe | 73 |
| HWL | M | 19 | 104 | Left frontal lobe | 80 |
| ZHX | M | 31 | 87 | Left medial temporal lobe | <51 |
| WXY | M | 24 | 108 | Left temporal lobe | 75 |
| LQM | F | 23 | 96 | Right temporal insular cortex | 81 |
| CLF | F | 21 | 94 | Bilateral medial temporo-occipital junction regions | 52 |
| HLL | F | 26 | 95 | Right temporo-occipital junction region | 70 |
| DHZ | F | 16 | 83 | Right limbic system | 77 |
| CYY | M | 35 | 83 | Right temporal lobe | 51 |
| HLY | M | 26 | 107 | Bilateral limbic system | 95 |
| YJH | M | 22 | 92 | Right temporo-insular region | 63 |
| GKS | M | 30 | 88 | Bilateral medial temporal lobes, right hippocampus, and right insula | <51 |
| ZH | M | 21 | 98 | Left temporo-insular lobe, orbital frontal gyrus, precentral sulcus, and parietal lobe. | 81 |
| KYP | M | 27 | 98 | Medial temporal lobe combined with insula, opercula, and orbitofrontal region | 73 |
| LYT | M | 18 | 91 | Right temporo-parietal junction region | 89 |
| LC | M | 36 | 93 | Left posterior temporal lobe | 84 |
| ZJJ | M | 21 | 93 | Right hippocampus, right insula, right central operculum | 108 |
| ZXF | M | 32 | 124 | Bilateral parieto-occipital regions | 120 |
| ZJH | M | 27 | 84 | Right posterior temporal region and temporo-occipital junction cortex | 52 |
| LHD | M | 33 | - | Left temporo-insular lobe | - |
| C | M | 23 | 82 | Right anterior temporal lobe | 66 |
| YLL | F | 33 | 75 | Right limbic system | 68 |
| WTT | F | 24 | - | Right temporal insular cortex | - |
| MAY | M | 40 | - | Right insula, right posterior orbital region and right medial temporal lobe | - |
| LHM | M | 25 | 97 | Left medial temporal lobe and insula | 54 |
| LSW | M | 21 | 82 | Left prefrontal lobe | 55 |
| CCL | M | 22 | - | Left temporo-insular lobe | - |
| LDN | F | 21 | 109 | Left temporal lobe | 103 |
| pat_02651_1127 | F | 27 | - | Left temporal lobe | - |
| pat_02660_1136 | F | 34 | - | Bilateral temporo-mesial | - |
| pat_02680_1158 | F | 52 | - | Left temporal lobe | - |
| pat_02689_1168 | F | 31 | 75 | Right temporo-parietal | - |
| pat_02711_1193 | F | 18 | 118 | Right temporal lobe | - |
| pat_02718_1201 | F | 26 | - | Left temporal lobe | - |
| pat_02985_1405 | M | 31 | - | undetermined focus | - |
| pat_03012_1444 | M | 26 | - | Temporo-polar | - |
| pat_03046_1482 | F | 39 | 93 | Right temporal | - |
| pat_03083_1527 | M | 25 | 101 | Bilateral temporo-basal | - |
| pat_03092_1538 | F | 35 | 81 | Bilateral temporo-mesial | - |
| pat_03105_1551 | F | 32 | - | Bilateral temporo-mesial | - |
| pat_03128_1591 | F | 28 | 99 | Bilateral temporo-mesial | - |
| pat_03138_1601 | F | 43 | - | Left temporo-mesial | - |
| pat_03146_1608 | M | 37 | 88 | Bilateral temporo-mesial | - |
| pat_03174_1634 | M | 25 | - | Polar and baso-medial region of the right temporal lobe | - |
| pat_03249_1711 | F | 52 | - | Left temporo-mesial | - |
| pat_03266_1729 | M | 39 | - | Bilateral temporo-mesial | - |
| pat_03300_1766 | F | 25 | - | Right temporo-mesial | - |

**Supplementary Table 1. Patient demographic and clinical information.** Details about patients’ gender, age, Intelligence Quotient (IQ), Seizure Onset Zone (SOZ), and Memory Quotient (MQ) are provided.

**Supplementary Table 2. Electrode coordinates in MNI space and Freesurfer labels for all subjects and electrodes**.

| ROI | CS++ | CS+- | CS-- |
| --- | --- | --- | --- |
| AMY | 38.41 ± 8.01 | 37.18 ± 8.84 | 37.78 ± 9.31 |
| HPC | 37.44 ± 8.32 | 36.53 ± 8.86 | 36.47 ± 8.72 |
| OFC | 39.17 ± 9.55 | 40.33 ± 9.57 | 39.83 ± 9.75 |
| lPFC | 40.78 ± 8.72 | 40.74 ± 9.67 | 41.26 ± 9.43 |
| TMP | 26.54 ±12.72 | 27.89 ± 12.41 | 26.85 ± 12.43 |

**Supplementary Table 3: Number of excluded trials after artifact rejection in every ROI across all task stages.** Please Note that since some of the RSA analyses rely on the comparison of many items of the same type (e.g., correlations are computed between all pairs of CS+ items during a specific task stage), the statistical power of these analyses is much higher. The number of trials in the table applies to the power analysis (Figure 2) and to the trial-level RSA analysis (Figure 3F and 4D).

**Supplementary Video 1. AMY and HPC electrodes visualization.** Visualization of Freesurfer average brain surfaces corresponding to the AMY (red) and the HPC (blue), in MNI coordinates. Electrodes localized within these regions and included in our analyses are displayed in red (AMY) and blue (HPC).

**Supplementary References**

Baas, J. M., Nugent, M., Lissek, S., Pine, D. S., & Grillon, C. (2004). Fear conditioning in virtual reality contexts: A new tool for the study of anxiety. *Biological Psychiatry*, *55*(11), 1056–1060. https://doi.org/10.1016/j.biopsych.2004.02.024

Balooch, S. B., Neumann, D. L., & Boschen, M. J. (2012). Extinction treatment in multiple contexts attenuates ABC renewal in humans. *Behaviour Research and Therapy*, *50*(10), 604–609. https://doi.org/10.1016/j.brat.2012.06.003

Bandarian Balooch, S., & Neumann, D. L. (2011). Effects of multiple contexts and context similarity on the renewal of extinguished conditioned behaviour in an ABA design with humans. *Learning and Motivation*, *42*(1), 53–63. https://doi.org/10.1016/j.lmot.2010.08.008

Battaglia, S., Garofalo, S., & di Pellegrino, G. (2018). Context-dependent extinction of threat memories: Influences of healthy aging. *Scientific Reports*, *8*(1), 12592. https://doi.org/10.1038/s41598-018-31000-9

Bernabei, J. M., Li, A., Revell, A. Y., Smith, R. J., Gunnarsdottir, K. M., Ong, I. Z., Davis, K. A., Sinha, N., Sarma, S., & Litt, B. (2023). Quantitative approaches to guide epilepsy surgery from intracranial EEG. *Brain*, *146*(6), 2248–2258. https://doi.org/10.1093/brain/awad007

Boddez, Y., Baeyens, F., Luyten, L., Vansteenwegen, D., Hermans, D., & Beckers, T. (2013). Rating data are underrated: Validity of US expectancy in human fear conditioning. *Journal of Behavior Therapy and Experimental Psychiatry*, *44*(2), 201–206. https://doi.org/10.1016/j.jbtep.2012.08.003

Bouyeure, A., Pacheco, D., Fellner, M.-C., Jacob, G., Kobelt, M., Rose, J., & Axmacher, N. (2024). Distinct representational properties of cues and contexts shape fear learning and extinction. *bioRxiv*, 2024.12.16.628638. https://doi.org/10.1101/2024.12.16.628638

Chen, S., Tan, Z., Xia, W., Gomes, C. A., Zhang, X., Zhou, W., Liang, S., Axmacher, N., & Wang, L. (2021). Theta oscillations synchronize human medial prefrontal cortex and amygdala during fear learning. *Science Advances*, *7*(34), eabf4198. https://doi.org/10.1126/sciadv.abf4198

Corcoran, K. A., & Maren, S. (2001). Hippocampal Inactivation Disrupts Contextual Retrieval of Fear Memory after Extinction. *The Journal of Neuroscience*, *21*(5), 1720. https://doi.org/10.1523/JNEUROSCI.21-05-01720.2001

Corcoran, K. A., & Maren, S. (2004). Factors regulating the effects of hippocampal inactivation on renewal of conditional fear after extinction. *Learning & Memory*, *11*(5), 598–603.

Desikan, R. S., Ségonne, F., Fischl, B., Quinn, B. T., Dickerson, B. C., Blacker, D., Buckner, R. L., Dale, A. M., Maguire, R. P., Hyman, B. T., Albert, M. S., & Killiany, R. J. (2006). An automated labeling system for subdividing the human cerebral cortex on MRI scans into gyral based regions of interest. *NeuroImage*, *31*(3), 968–980. https://doi.org/10.1016/j.neuroimage.2006.01.021

Dimakopoulos, V., Gotman, J., Stacey, W., von Ellenrieder, N., Jacobs, J., Papadelis, C., Cimbalnik, J., Worrell, G., Sperling, M. R., Zijlmans, M., Imbach, L., Frauscher, B., & Sarnthein, J. (2022). Protocol for multicentre comparison of interictal high-frequency oscillations as a predictor of seizure freedom. *Brain Communications*, *4*(3), fcac151. https://doi.org/10.1093/braincomms/fcac151

Dunsmoor, J. E., Ahs, F., Zielinski, D. J., & LaBar, K. S. (2014). Extinction in multiple virtual reality contexts diminishes fear reinstatement in humans. *Extinction*, *113*, 157–164. https://doi.org/10.1016/j.nlm.2014.02.010

Dunsmoor, J. E., Bandettini, P. A., & Knight, D. C. (2007). Impact of continuous versus intermittent CS-UCS pairing on human brain activation during Pavlovian fear conditioning. *Behavioral Neuroscience*, *121*(4), 635.

Dunsmoor, J. E., Martin, A., & LaBar, K. S. (2012). Role of conceptual knowledge in learning and retention of conditioned fear. *Biological Psychology*, *89*(2), 300–305. https://doi.org/10.1016/j.biopsycho.2011.11.002

Dunsmoor, J. E., & Murphy, G. L. (2014). Stimulus Typicality Determines How Broadly Fear Is Generalized. *Psychological Science*, *25*(9), 1816–1821. https://doi.org/10.1177/0956797614535401

Effting, M., & Kindt, M. (2007). Contextual control of human fear associations in a renewal paradigm. *Behaviour Research and Therapy*, *45*(9), 2002–2018. https://doi.org/10.1016/j.brat.2007.02.011

Fell, J., & Axmacher, N. (2011). The role of phase synchronization in memory processes. *Nature Reviews. Neuroscience*, *12*(2), 105–118. https://doi.org/10.1038/nrn2979

Garcia, J., & Koelling, R. A. (1966). Relation of cue to consequence in avoidance learning. *Psychonomic Science*, *4*(3), 123–124. https://doi.org/10.3758/BF03342209

Glenn, C. R., Klein, D. N., Lissek, S., Britton, J. C., Pine, D. S., & Hajcak, G. (2012). The development of fear learning and generalization in 8–13 year-olds. *Developmental Psychobiology*, *54*(7), 675–684. https://doi.org/10.1002/dev.20616

Glenn, C. R., Lieberman, L., & Hajcak, G. (2012). Comparing electric shock and a fearful screaming face as unconditioned stimuli for fear learning. *International Journal of Psychophysiology*, *86*(3), 214–219. https://doi.org/10.1016/j.ijpsycho.2012.09.006

Glotzbach-Schoon, E., Tadda, R., Andreatta, M., Tröger, C., Ewald, H., Grillon, C., Pauli, P., & Mühlberger, A. (2013). Enhanced discrimination between threatening and safe contexts in high-anxious individuals. *Biological Psychology*, *93*(1), 159–166. https://doi.org/10.1016/j.biopsycho.2013.01.011

Hamm, A. O., Vaitl, D., & Lang, P. J. (1989). Fear conditioning, meaning, and belongingness: A selective association analysis. *Journal of Abnormal Psychology*, *98*(4), 395–406. https://doi.org/10.1037/0021-843X.98.4.395

Henin, S., Shankar, A., Borges, H., Flinker, A., Doyle, W., Friedman, D., Devinsky, O., Buzsáki, G., & Liu, A. (2021). Spatiotemporal dynamics between interictal epileptiform discharges and ripples during associative memory processing. *Brain*, *144*(5), 1590–1602. https://doi.org/10.1093/brain/awab044

Hermann, A., Stark, R., Milad, M., & Merz, C. (2016). Renewal of conditioned fear in a novel context is associated with hippocampal activation and connectivity. *Social Cognitive and Affective Neuroscience*, *11*(9), 1411–1421.

Imperatori, L. S., Betta, M., Cecchetti, L., Canales-Johnson, A., Ricciardi, E., Siclari, F., Pietrini, P., Chennu, S., & Bernardi, G. (2019). EEG functional connectivity metrics wPLI and wSMI account for distinct types of brain functional interactions. *Scientific Reports*, *9*(1), 8894. https://doi.org/10.1038/s41598-019-45289-7

Kalisch, R., Korenfeld, E., Stephan, K. E., Weiskopf, N., Seymour, B., & Dolan, R. J. (2006). Context-Dependent Human Extinction Memory Is Mediated by a Ventromedial Prefrontal and Hippocampal Network. *The Journal of Neuroscience*, *26*(37), 9503. https://doi.org/10.1523/JNEUROSCI.2021-06.2006

Krisch, K. A., Bandarian-Balooch, S., & Neumann, D. L. (2018). Effects of extended extinction and multiple extinction contexts on ABA renewal. *Learning and Motivation*, *63*, 1–10. https://doi.org/10.1016/j.lmot.2017.11.001

Lachaux, J.-P., Rodriguez, E., Martinerie, J., & Varela, F. J. (1999). Measuring phase synchrony in brain signals. *Human Brain Mapping*, *8*(4), 194–208. https://doi.org/10.1002/(SICI)1097-0193(1999)8:4<194::AID-HBM4>3.0.CO;2-C

Lonsdorf, T. B., Menz, M. M., Andreatta, M., Fullana, M. A., Golkar, A., Haaker, J., Heitland, I., Hermann, A., Kuhn, M., Kruse, O., Meir Drexler, S., Meulders, A., Nees, F., Pittig, A., Richter, J., Römer, S., Shiban, Y., Schmitz, A., Straube, B., … Merz, C. J. (2017). Don’t fear ‘fear conditioning’: Methodological considerations for the design and analysis of studies on human fear acquisition, extinction, and return of fear. *Neuroscience & Biobehavioral Reviews*, *77*, 247–285. https://doi.org/10.1016/j.neubiorev.2017.02.026

Michelmann, S., Bowman, H., & Hanslmayr, S. (2016). The Temporal Signature of Memories: Identification of a General Mechanism for Dynamic Memory Replay in Humans. *PLOS Biology*, *14*(8), e1002528. https://doi.org/10.1371/journal.pbio.1002528

Milad, M. R., Orr, S. P., Pitman, R. K., & Rauch, S. L. (2005). Context modulation of memory for fear extinction in humans. *Psychophysiology*, *42*(4), 456–464.

Milad, M. R., Wright, C. I., Orr, S. P., Pitman, R. K., Quirk, G. J., & Rauch, S. L. (2007). Recall of Fear Extinction in Humans Activates the Ventromedial Prefrontal Cortex and Hippocampus in Concert. *Biological Psychiatry*, *62*(5), 446–454. https://doi.org/10.1016/j.biopsych.2006.10.011

Mineka, S., & Öhman, A. (2002). Phobias and preparedness: The selective, automatic, and encapsulated nature of fear. *Biological Psychiatry*, *52*(10), 927–937. https://doi.org/10.1016/S0006-3223(02)01669-4

Mormann, F., Lehnertz, K., David, P., & E. Elger, C. (2000). Mean phase coherence as a measure for phase synchronization and its application to the EEG of epilepsy patients. *Physica D: Nonlinear Phenomena*, *144*(3), 358–369. https://doi.org/10.1016/S0167-2789(00)00087-7

Neumann, D. L., Lipp, O. V., & Cory, S. E. (2007). Conducting extinction in multiple contexts does not necessarily attenuate the renewal of shock expectancy in a fear-conditioning procedure with humans. *Behaviour Research and Therapy*, *45*(2), 385–394. https://doi.org/10.1016/j.brat.2006.02.001

Ney, L. J., O’Donohue, M. P., Lowe, B. G., & Lipp, O. V. (2022). Angry and fearful compared to happy or neutral faces as conditional stimuli in human fear conditioning: A systematic review and meta-analysis. *Neuroscience & Biobehavioral Reviews*, *139*, 104756. https://doi.org/10.1016/j.neubiorev.2022.104756

Nolte, G., Ziehe, A., Nikulin, V. V., Schlögl, A., Krämer, N., Brismar, T., & Müller, K.-R. (2008). Robustly estimating the flow direction of information in complex physical systems. *Physical Review Letters*, *100*(23), 234101.

Öhman, A., & Öst, L.-G. (1985). Animal and social phobias: Biological constraints on learned fear responses. *Theoretical Issues in Behavior Therapy*, 123–175.

Pacheco Estefan, D., Sánchez-Fibla, M., Duff, A., Principe, A., Rocamora, R., Zhang, H., Axmacher, N., & Verschure, P. F. M. J. (2019). Coordinated representational reinstatement in the human hippocampus and lateral temporal cortex during episodic memory retrieval. *Nature Communications*, *10*(1), 1–13.

Schmitz, A., & Grillon, C. (2012). Assessing fear and anxiety in humans using the threat of predictable and unpredictable aversive events (the NPU-threat test). *Nature Protocols*, *7*(3), 527–532.

Stam, C. J., Nolte, G., & Daffertshofer, A. (2007). Phase lag index: Assessment of functional connectivity from multi channel EEG and MEG with diminished bias from common sources. *Human Brain Mapping*, *28*(11), 1178–1193.

Vansteenwegen, D., Vervliet, B., Iberico, C., Baeyens, F., Van den Bergh, O., & Hermans, D. (2007). The repeated confrontation with videotapes of spiders in multiple contexts attenuates renewal of fear in spider-anxious students. *Behaviour Research and Therapy*, *45*(6), 1169–1179. https://doi.org/10.1016/j.brat.2006.08.023

Varela, F., Lachaux, J.-P., Rodriguez, E., & Martinerie, J. (2001). The brainweb: Phase synchronization and large-scale integration. *Nature Reviews Neuroscience*, *2*(4), 229–239.

Vinck, M., Oostenveld, R., van Wingerden, M., Battaglia, F., & Pennartz, C. M. A. (2011). An improved index of phase-synchronization for electrophysiological data in the presence of volume-conduction, noise and sample-size bias. *NeuroImage*, *55*(4), 1548–1565. https://doi.org/10.1016/j.neuroimage.2011.01.055

Visser, R. M., Scholte, H. S., Beemsterboer, T., & Kindt, M. (2013). Neural pattern similarity predicts long-term fear memory. *Nature Neuroscience*, *16*(4), 388–390. https://doi.org/10.1038/nn.3345

Wehrli, J. M., Xia, Y., Gerster, S., & Bach, D. R. (2022). Measuring human trace fear conditioning. *Psychophysiology*, *59*(12), e14119. https://doi.org/10.1111/psyp.14119

Xia, Y., Wehrli, J., Gerster, S., Kroes, M., Houtekamer, M., & Bach, D. R. (2023). Measuring human context fear conditioning and retention after consolidation. *Learning & Memory*, *30*(7), 139–150.
